# Sex of donor cell and reprogramming conditions predict the extent and nature of imprinting defects in mouse iPSCs

**DOI:** 10.1101/2020.11.20.370973

**Authors:** Maria Arez, Melanie Eckersley-Maslin, Tajda Klobučar, João von Gilsa Lopes, Felix Krueger, Ana Cláudia Raposo, Anne-Valerie Gendrel, Bruno Bernardes de Jesus, Simão Teixeira da Rocha

## Abstract

Reprogramming of somatic cells into induced Pluripotent Stem Cells (iPSCs) is a major leap towards personalized approaches to disease modelling and cell-replacement therapies. However, we still lack the ability to fully control the epigenetic status of iPSCs, which is a major hurdle for their downstream applications. A sensible indicator for epigenetic fidelity is genomic imprinting, a phenomenon dependent on DNA methylation, which is frequently perturbed in iPSCs by yet unidentified reasons. By using a secondary reprogramming system with murine hybrid donor cells, we conducted a thorough imprinting analysis using IMPLICON in multiple female and male iPSCs generated under different culture conditions. Our results show that imprinting defects are remarkably common in mouse iPSCs causing dysregulation of the typical monoallelic expression of imprinted genes. Interestingly, the nature of imprinting defects depends on the sex of the donor cell and their respective response to culture conditions. Under serum-free conditions, male iPSCs show global hypomethylation at imprinted regions, whereas in serum conditions show focal hypermethylation at specific loci. In contrast, female iPSCs always exhibit hypomethylation defects regardless of culture conditions. These imprinting defects are more severe than the global changes in DNA methylation, highlighting the sensitivity of imprinting loci to current iPSC generation protocols. Our results reveal clear predictors underlying different types of imprinting defects in mouse iPSCs. This knowledge is essential to devise novel reprogramming strategies aiming at generating epigenetically faithful iPSCs.

## INTRODUCTION

The seminal studies from Takahashi and Yamanaka over a decade ago demonstrated the possibility to revert a somatic cell to a stem-like state through the induction of a defined set of transcription factors (Takahashi and Yamanaka, 2006). This opened the prospect to reprogram patient-derived cells, helping to elucidate novel pathological mechanisms underlying human diseases and to reveal new therapeutic drugs. Furthermore, clinical trials involving iPSCs in cell-replacement therapies have already been initiated (Mandai et al., 2017; Shi et al., 2017).

Despite being a revolutionary technology, several challenges persist that limit the current range of iPSC applications. First, iPSC reprogramming remains a highly inefficient process with most of the cells either failing or achieving only partial reprogramming (Buganim et al., 2013; Schiebinger et al., 2019; Tonge et al., 2014; Yamanaka, 2009). Second, correctly reprogrammed iPSCs exhibit very heterogeneous responses to specific differentiation cues, which compromise their use in disease modelling and clinical applications (Bar-Nur et al., 2011; Kim et al., 2011; Nishizawa et al., 2016). Lack of consistency between different iPSCs can be attributed to the natural genetic variability among individual donors (Kajiwara et al., 2012), as well as recurrent genetic aberrations in iPSCs (Avior et al., 2019; Ben-David et al., 2014; Gore et al., 2011). However, the persistence of such heterogeneous behaviour in isogenic iPSC lines suggests also a role of epigenetic variability in this phenomenon (Bock et al., 2011; Ma et al., 2014; Nazor et al., 2012; Nishizawa et al., 2016).

Epigenetic disparity, mostly studied at the DNA methylation level, can originate from: (1) failure to fully reset the somatic memory of the donor cell (Bar-Nur et al., 2011; Kim et al., 2011; Lister et al., 2011; Ohi et al., 2011; Polo et al., 2010); (2) acquisition of aberrations during the reprogramming process (Huang et al., 2014; Koyanagi-Aoi et al., 2013; Lister et al., 2011; Ma et al., 2014; Nazor et al., 2012; Ohi et al., 2011; Ruiz et al., 2012; Stadtfeld et al., 2010); (3) epigenetic adaptation to long-term *in vitro* culturing (Mekhoubad et al., 2012; Vallot et al., 2015).The latter two likely affect processes that rely on strict maintenance of DNA methylation such as genomic imprinting (Stadtfeld et al., 2010; Sun et al., 2012; Takikawa et al., 2013; Yagi et al., 2019), which serves as a good readout to study epigenetic fidelity in iPSCs.

Genomic imprinting is a parent-of-origin specific epigenetic mechanism that controls the monoallelic expression of ~150 genes in the mammalian genome. Unlike the majority of genes, imprinted genes are biased or exclusively expressed from the maternal or paternal chromosomes. These genes are important regulators in prenatal growth and development, as well as, in postnatal brain functions and metabolic pathways [reviewed in (Tucci et al., 2019)]. The dysregulation of imprinted genes has been associated with several developmental and behavioural disorders, such as Beckwith-Weidemann, Angelman and Prader-Willi syndromes (Cassidy et al., 2012; Maranga et al., 2020; Wang et al., 2019). Most imprinted genes are located in close proximity in defined genomic loci, called imprinted clusters. Mammalian genomes present around 25 imprinted clusters, that contain a *cis*-acting regulatory element, known as imprinting control region (ICR), which co-regulates imprinting expression of multiple genes. This region is dense in CpG dinucleotides and displays opposite DNA methylation patterns between the paternally and maternally inherited alleles. This methylation pattern is set up during gametogenesis and stably maintained in somatic cells, with 22 of the ICRs being methylated on the maternal allele and 3 on the paternal allele in the mouse genome [reviewed in (Tucci et al., 2019)]. Disturbances in the parental allele-specific methylation at ICRs perturb monoallelic expression and are thus one of the main causes of imprinted disorders (Soellner et al., 2017). Therefore, it is of utmost importance that proper parental-specific methylation at ICRs is maintained in iPSCs.

Previous studies have revealed imprinting defects in both mouse and human iPSCs (Bar et al., 2017; Ma et al., 2014; Nazor et al., 2012; Stadtfeld et al., 2010; Sun et al., 2012; Takikawa et al., 2013; Yagi et al., 2019). Importantly, these errors persist and are never rescued upon differentiation (Bar et al., 2017; Nazor et al., 2012), which is troublesome for iPSC applications in translational and clinical research. The first reported imprinting defect was the aberrant hypermethylation of the maternal allele in the *Dlk1-Dio3* cluster leading to silencing of maternally expressed non-coding RNAs in mouse iPSCs (Stadtfeld et al., 2010). This was shown to compromise their pluripotent properties (Carey et al., 2011; Liu et al., 2010; Stadtfeld et al., 2010). Interestingly, defects in *Dlk1-Dio3* loci could be corrected by using ascorbic acid (Stadtfeld et al., 2012), presumably due to an increased demethylating activity of the TET dioxygenase enzymes (Blaschke et al., 2013). Later studies showed that the repertoire of imprinting defects extends to other imprinted loci in mouse iPSCs (Sun et al., 2012; Takikawa et al., 2013; Yagi et al., 2019). Interestingly, the extent and nature (hypermethylation versus hypomethylation) of these defects varied significantly between studies (Table S1). Whether these contrasting results are caused by differences in the sex of donor cells and/or the reprogramming protocols used, or by limitations of the systems chosen for imprinting analyses is unknown (Table S1). Several shortcomings can be attributed to those studies which prevent direct comparison: (1) unavailability of single nucleotide polymorphisms (SNPs) to distinguish the two parental alleles for imprinting analysis; (2) low number of iPSC lines studied; (3) lack of information or use of iPSCs of only one sex; (4) use of a single culture condition for reprogramming; (5) limited number of imprinted regions assessed.

Here, we present a systematic analysis of imprinting defects in isogenic murine female and male hybrid iPSCs in serum-free and serum-based culture conditions using an ultra-deep approach to screen imprinting methylation. Our results show that virtually all generated iPSCs have imprinting errors, highlighting, unequivocally, the persistence of these epigenetic errors in iPSCs. Moreover, the nature of these errors are dictated by sex of the donor cell and medium conditions. Our results give important perspectives towards the design of novel reprogramming strategies to generate iPSCs with a stable epigenome.

## RESULTS

### Generation and characterization of F1 hybrid iPSCs in serum-free conditions

To systematically address the imprinting status in mouse iPSCs, we generated male and female isogenic hybrid iPSC lines from mouse embryonic fibroblasts (MEFs) derived from a cross between a female “reprogrammable” i4F mouse carrying a doxycycline-inducible polycistronic Yamanaka cassette (on a C57BL/6J background; named i4F-BL6 herein) (Abad et al., 2013; Bernardes de Jesus et al., 2018) and a male from the *Mus musculus castaneus* (CAST/Ei - named CAST herein) strain. The use of F1 hybrid cells from genetically distant mouse strains allows the distinction of parental alleles based on SNPs present at both ICRs and genes (Strogantsev et al., 2015; Xie et al., 2012). Unfortunately, we were unable to obtain MEFs from the reciprocal cross within the time-frame of this work. For this reason, we focused our analysis on loci for which imprinting has previously been shown not to be perturbed by the direction of the cross either in tissues or in mouse iPSCs (Klobucar et al., 2020; Sun et al., 2012; Xie et al., 2012).

Both female and male MEFs were first reprogrammed in serum-free conditions using medium supplemented with Knock-out serum replacement (KSR) in the presence of doxycycline (DOX) for 12 days (Fig. 1A) as previously described (Bernardes de Jesus et al., 2018). After 12 days of DOX induction, we picked 6 independent colonies with the typical dome-shaped morphology from each sex. The colonies were maintained in the same medium without DOX until around day 50 to ensure the generation of totally reprogramed iPSCs (Fig. 1A). All the analyses were performed at this stage and in the subsequent two-to-three passages.

**Fig. 1.**
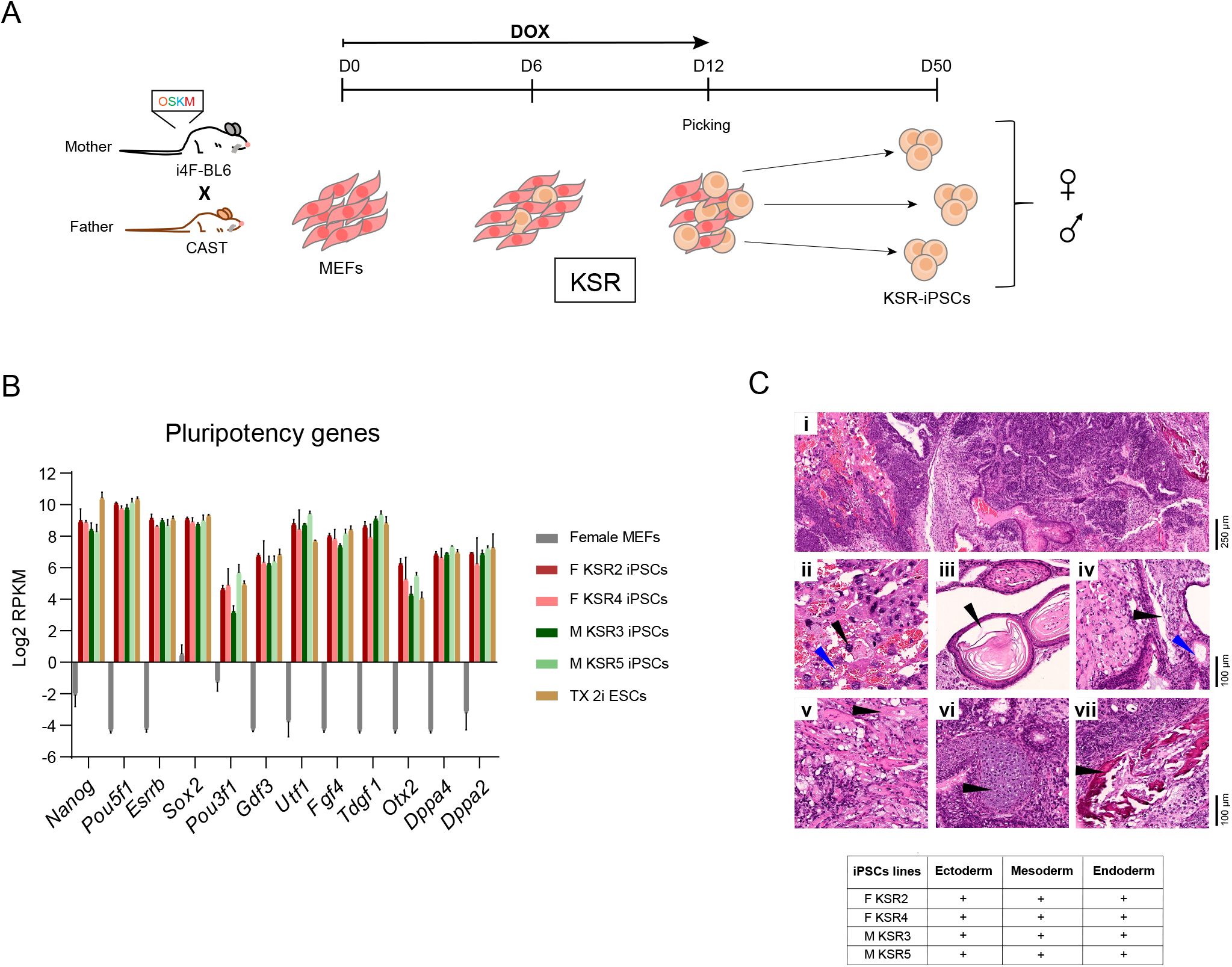
Generation of KSR-derived F1 hybrid iPSCs. A. Schematic representation of the reprogramming protocol; briefly, a transgenic “reprogrammable” female mouse on a C57BL/6J genetic background (i4F-BL6) was crossed with a *Mus musculus castaneus* (CAST) male mouse to generate E13.5 F1 hybrid embryos to generate mouse embryonic fibroblasts (MEFs). MEFs were reprogrammed by induction of the polycistronic Yamanaka cassette (*Oct4*/*Sox2*/*Klf4*/*c-Myc* - OSKM) in the presence of doxycycline (DOX) for 12 days. Individual clones of mouse induced pluripotent stem cells reprogrammed in Knockout Serum Replacement medium (KSR-iPSCs) were picked at day 12 and expanded until approximately day 50. B. Expression analysis by RNAseq of a panel of pluripotent genes in female MEFs, female (F KSR2, F KSR3), male (M KSR3 and M KSR5) iPSCs and TX 2i ESCs. Graph shows the average Log2 Reads per kilobase per million mapped reads (RPKM) expression values ± Standard Deviation (SD) from biological triplicates of each sample. C. Table and representative H&E staining of teratomas after subcutaneous injection of 2 × 10^6^ cells into the flanks of NSG mice. iPSCs efficiently contribute to ectoderm, mesoderm, endoderm and occasionally trophectoderm. **i**, Low magnification of a mature teratoma, scale bar represents 250 μm. **ii**, Trophectoderm-derived trophoblast giant cells (black arrowhead), associated with large vascular spaces (blue arrowhead), characteristic of placental tissue. **iii**, Ectodermal components corresponding to squamous epithelium (black arrowhead). **iv**, Endodermal components corresponding to respiratory-type epithelium, including ciliated (black arrowhead), and mucin-producing goblet cells (blue arrowhead). **v, vi, vii**, Mesodermal components (black arrowhead) corresponding to muscle, cartilage, and bone, respectively; ii-vii scale bar represents 100 μm. Table summarizes the successful generation of teratomas with tissues from the three germ layers from F KSR2, F KSR4, M KSR3 and M KSR5 iPSCs.

To validate the newly KSR-derived iPSCs (KSR-iPSCs), we screened for the expression of three pluripotent markers (*Pou5f1*, *Nanog* and *Esrrb*). All KSR-iPSCs, but not the parental MEFs, expressed the markers at similar levels to mouse embryonic stem cells (ESCs) cultured in serum (JM8.F6 male ESCs) or 2i (MEK and GSK3 inhibition) conditions (TX1072 female ESCs from a BL6 female x CAST male cross – TX 2i ESCs) (Schulz et al., 2014) (Fig. S1A). We strengthened our transcriptional characterization by performing RNA sequencing (RNA-seq) in three biological replicates for two female (F KSR2 and F KSR4) and two male (M KSR3 and M KSR5) iPSC lines, TX 2i ESCs, as well as female MEFs. Clustering analysis based on expression profile demarcates differences between cell types (MEFs versus iPSCs/ESCs) and culture conditions (KSR versus 2i) (Fig. S1B). Analysis of a selected set of pluripotency genes showed consistent expression in iPSCs and TX 2i ESCs, but not in female MEFs (Fig. 1B). This was confirmed at the protein level by immunofluorescence (IF) for SSEA-1, OCT4 and NANOG (Fig. S1C). RNAseq clustering analysis did not separate iPSCs according to the sex, but differential expression analysis revealed 173 downregulated and 413 upregulated genes in female versus male iPSCs (Fig. S1D; Table S2). Most (246 out of 413) of the upregulated genes are mapped on the X chromosome and no X-linked genes were downregulated, consistent with the fact that female iPSCs/ESCs have two active X chromosomes (Fig. S1D). When female and male KSR-iPSC lines were subcutaneously injected into immune-deficient NOD SCID γ (NSG) mice (Shultz et al., 2007), they all formed teratomas composed by cells belonging to the three germ layers (endoderm, mesoderm and ectoderm) and the occasional presence of trophectoderm-derived trophoblast giant cells, confirming their full differentiation potential (Fig. 1C). All together, these results show that we successfully generated multiple pluripotent female and male hybrid KSR-iPSCs.

### Female and male KSR-iPSCs show loss of methylation at imprinted regions

To screen for imprinting methylation fidelity in KSR-iPSCs, we employed allele-specific IMPLICON previously validated on F1 tissue samples from reciprocal BL6 x CAST crosses (Klobucar et al., 2020). This method combines bisulfite treatment of genomic DNA with amplicon high-throughput sequencing with a de-duplication step to generate base-resolution datasets with high coverage and allelic discrimination of the original DNA molecules. We focused on 8 imprinted clusters together with 2 unmethylated and 1 methylated control regions. As expected, no differences between the two parental alleles were observed for both unmethylated (*Sox2* and *Klf4* genes) and methylated controls (*Prickle1* gene) in female and male MEFs and respective KSR-iPSCs, as well as, for the TX 2i ESCs (Fig. S2A; Table S3). For *Prickle1*, a slight drop in DNA methylation levels was observed for female KSR-iPSCs (~50-70%) in both alleles compared to parental MEFs (>90%), while it was completely lost in the TX 2i ESC line (<10%), likely to be caused by 2i-induced demethylation (Ficz et al., 2013; Habibi et al., 2013; Leitch et al., 2013; von Meyenn et al., 2016).

For the 8 imprinted regions analysed, the expected allele-specific methylation pattern at ICRs was always observed for both female and male MEFs, with six of the regions displaying methylation on the maternal allele (PWS/AS*, Peg3, Gnas, Commd1-Zrsr1*, *Mcts2-H13*, *Kcqn1-Kcnq1ot1*) and two regions on the paternal allele (*Igf2-H19* and *Dlk1-Dio3*) (Fig. 2A; Fig. S2B; Table S3). This allele-specific pattern was mostly erased in the TX 2i ESCs, confirming that the 2i-induced demethylation does not spare imprinted loci as previously reported (Choi et al., 2017b; Lee et al., 2018). Strikingly, the expected imprinting methylation status was no longer found in either female or male KSR-iPSCs. First, we observed that reduction in methylation from the methylated ICRs was a common trend to all KSR-iPSCs (Fig. 2A; Fig. S2B; Table S3), and affected both maternally and paternally methylated ICRs. Second, these hypomethylation defects were more pronounced in female versus male KSR-iPSCs. This is particularly noticeable looking at averaged methylation levels of female and male KSR-iPSCs versus the original MEFs (Fig. 2B). While all 8 imprinted regions displayed significant hypomethylation defects in all female KSR-iPSCs, this was only true for 6 imprinted regions in male KSR-iPSCs, being the PWS/AS and *Dlk1-Dio3* clusters the exceptions (Fig. 2B). Third, we also noticed considerable iPSC-to-iPSC variation in methylation levels at imprinted regions (Fig. 2A; Fig. S2B) matching previous observations (Sun et al., 2012; Takikawa et al., 2013; Yagi et al., 2019).

**Figure 2.**
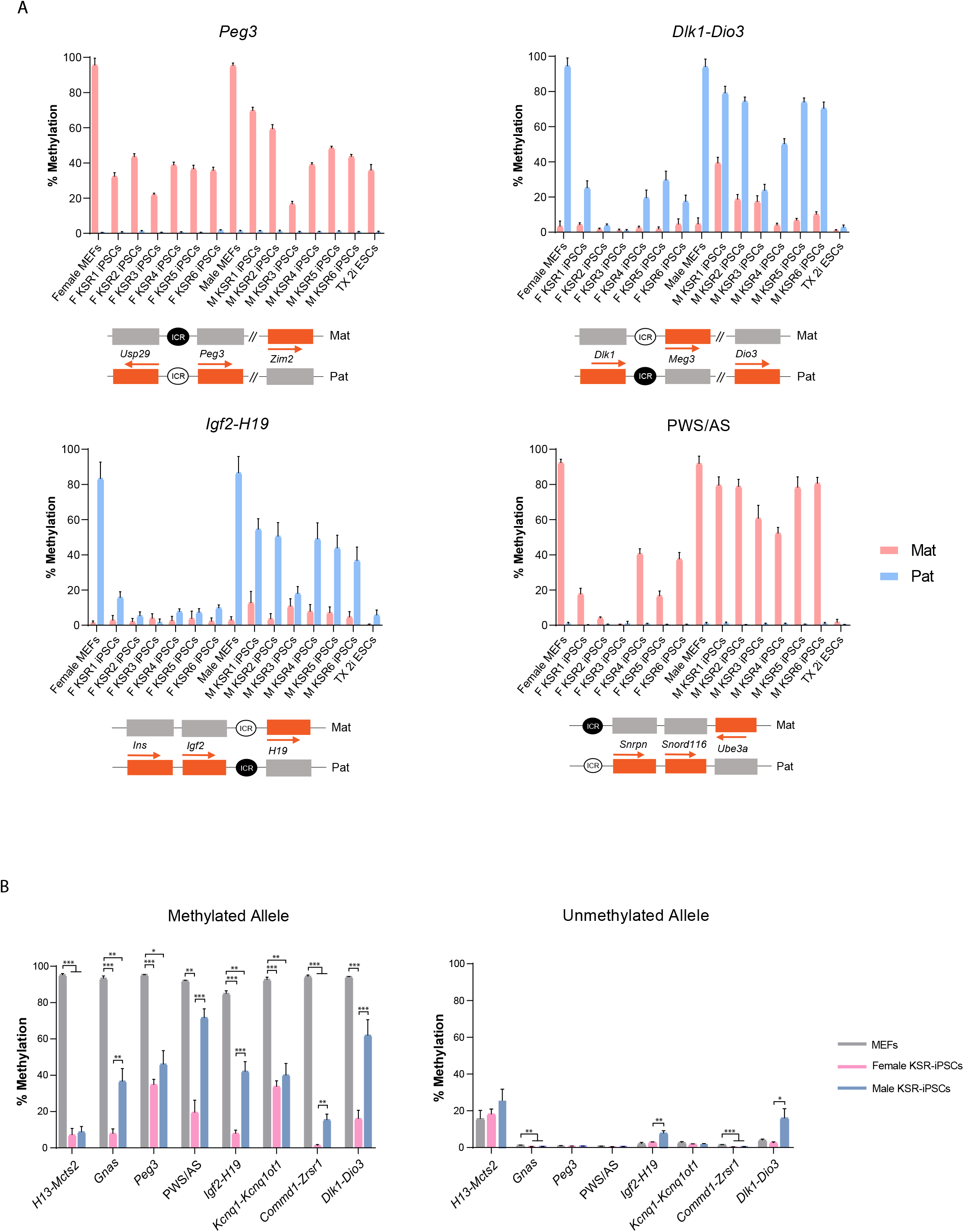
Hypomethylation defects in KSR-iPSCs. A. Methylation analysis of *Peg3*, *Dlk1-Dio3, Igf2-H19* and PWS/AS ICRs in female and male MEFs, female (F KSR1-F KSR6) and male (M KSR1-M KSR6) iPSCs and TX 2i ESCs; Each graph represents the mean percentage ± SD methylation levels measured at each CpG within different genomic regions per parental allele for each sample; Scheme on the bottom of each graph represents the normal methylation status of each ICR in the correspondent regions (white circle – unmethylated ICR; black circle – methylated ICR; Mat – maternal allele; Pat – paternal allele; orange rectangles –expressed genes; grey rectangles – silenced genes; regions are not drawn to scale). B. Average percentage of methylation at methylated and unmethylated ICRs in parental MEFs, female and male KSR-iPSCs; Graph represents the mean ± SEM methylation levels measured at each CpG within different genomic regions per parental allele for each group of samples. Statistically significant differences are indicated as * *p* < 0.05; ** *p* < 0.01; *** *p* < 0.001 (unpaired two-tailed Student’s *t*-test).

Hypomethylation defects ranged from a milder reduction in methylation (30-70%) to the complete loss of methylation (< 10%) at the methylated allele (Fig. 2A; Fig.S2B). Milder cases could either be explained by partial methylation loss at ICRs in all cells equally or by cellular heterogeneity within the same iPSC line, having some cells completely lost DNA methylation at ICRs, while others maintained it intact. Thanks to the single-molecule resolution of IMPLICON, we could see that the majority of these mild cases can indeed be explained by cellular heterogeneity. At the *Gnas* and *Kcnq1-Kcnq1ot1* loci, a partition between unmethylated and methylated reads could be seen for the normally methylated maternal ICRs (Fig. S2C), suggesting that some cells have lost DNA methylation at these elements, while others have retained it. Overall, our results show that both female and male KSR-iPSCs have substantial hypomethylation defects suggestive of a general tendency of imprinting erasure under these reprogramming conditions.

### Loss of imprinting methylation correlates with aberrations in parental allele-specific expression in KSR-iPSCs

We next examined whether ICR hypomethylation disrupts the normal allele-specific expression of imprinted genes by analysing our RNA-seq datasets taking advantage of the strain-specific SNPs. From a list of known murine imprinted genes (Tucci et al., 2019) with stable imprinting in BL6 x CAST reciprocal crosses, we shortlisted 17 imprinted genes (13 paternally and 4 maternally expressed) transcribed from a single allele in MEFs (ratio: > 90%:10%), expressed in all iPSC replicates (Log2 RPKM > 1) and with sufficient allelic resolution (normalized cumulative SNP-specific read counts > 5) in at least two of the three replicates (Fig. 3A; Table S4). Consistent with the widespread hypomethylation defects in KSR-iPSCs and TX 2i ESCs (Fig. 2A-B; Fig. S2B), we no longer found the expected parental allele-specific expression for the vast majority of imprinted genes (Fig. 3A). Most imprinted genes became biallelically expressed with three genes (e.g. *Meg3, Peg3* and *Plagl1*) even showing bias towards the originally silenced allele. The apparent switch of parental allele expression seen for the *Sgce*, *Peg10* and *Mest* genes (Fig. 3A) in the F KSR2 iPSC line was later confirmed to be caused by loss of the paternal-derived chromosome 6 in the majority of cells (Fig. S3A-B). Loss of monoallelic expression can be exemplified for the maternally *H19* expressed gene in female and male KSR-iPSCs, as well as in TX 2i ESCs (Fig. 3B). We also found that loss of monoallelic expression was dependent on the degree of the hypomethylation defect. For example, in the PWS/AS cluster, the paternally expressed *Snrpn* gene becomes biallelic in F KSR2 iPSCs and TX 2i ESCs due to loss of maternal ICR methylation, but remains monoallelically expressed in the barely affected M KSR5 iPSCs (Fig. 2A; Fig. 3B). Alterations in allele-specific expression were broadly consistent with the methylation changes at ICRs (Fig. 2A-B; Fig. 3A-B), revealing that both female and male KSR-iPSCs are incapable to maintain their imprinting expression.

**Figure 3.**
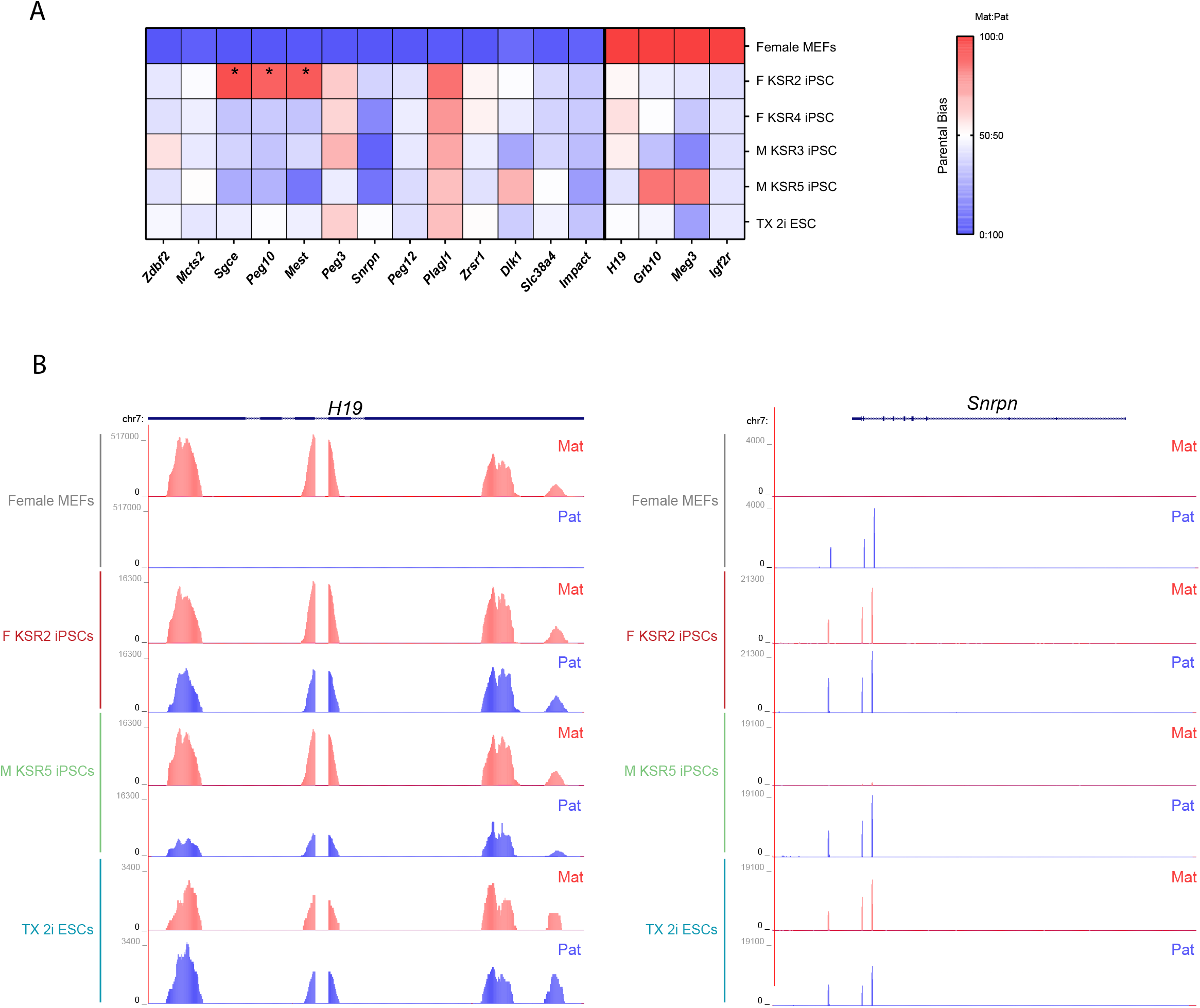
Dysregulation of imprinted expression in KSR-iPSCs. A. Heatmap representing the mean parental allele-specific expression biases of imprinted genes in the biological triplicates of female MEFs, female (F KSR2 and F KSR4) and male (M KSR3 and M KSR5) iPSCs and TX 2i ESCs. Asterisks signal genes exhibiting parental-specific expression biases induced by genetic alterations (see Fig. S3A). B. Genome browser plots showing allele-specific reads in the *H19* and *Snrpn* genes in female MEFs, F KSR2 and M KSR5 iPSCs and TX 2i ESCs. Maternal reads are shown in red and paternal reads in blue. Note that due to SNP distribution not all parts of the transcript can be assessed in an allele-specific manner.

### Imprinting defects in FBS-iPSCs depend on the sex of the donor cell

Given our findings of major hypomethylation defects at ICRs in KSR-iPSCs, we next asked whether switching to different culture conditions would change this phenotype. We applied the same reprogramming paradigm but using the classical ESC medium conditions based on fetal bovine serum (FBS) (Fig. 4A). We successfully generated 5 female and 5 male FBS-derived iPSCs (FBS-iPSCs) exhibiting normal ESC-like morphology, expressing pluripotent stem cell markers and forming well-differentiated teratomas (Fig. S4A-C). Allele-specific IMPLICON was then performed for the same 8 imprinted regions and methylated/unmethylated controls. No differences were observed for both the unmethylated (*Sox2* and *Klf4*) and methylated (*Prickle1*) control regions (Table S3). Female FBS-iPSCs exhibited lower levels of methylation in both alleles for *Prickle1* as previously seen in female KSR-iPSCs (Table S3). Strikingly, we found different imprinting outcomes in female versus male FBS-iPSCs. Similar to KSR-iPSCs (Fig. 2A-B), female FBS-iPSCs showed a strong tendency to demethylate the methylated ICRs (Fig. 4B; Fig. S4D-E; Table S3), resulting in biallelic expression of imprinted genes, as illustrated for the *H19* and *Snrpn* genes (Fig. 4C). In contrast, male FBS-iPSCs preserved the typical imprinting methylation status for 6 of the 8 imprinted regions showing correct monoallelic expression of imprinted genes (Fig. 4C), while two other regions exhibit signs of hypermethylation of the unmethylated ICR (Fig. 4B; Fig. S4D-E). The *Igf2-H19* cluster was one of the two regions with a tendency to gain methylation at the unmethylated allele (Fig. S4D-E), however, this did not seem to perturb the monoallelic expression of *H19* (Fig. 4C). A stronger gain of methylation was seen in the *Dlk1-Dio3* region, where 4 out of 5 iPSC clones showed equivalent high methylation levels in both parental alleles (Fig. 4B). Our results revealed marked differences in imprinting fidelity between female and male iPSCs reprogrammed under standard serum conditions. We could detect sex-specific differences in the nature of imprinting defects with hypomethylation linked to female, and hypermethylation defects at specific loci linked to male FBS-iPSCs.

**Figure 4.**
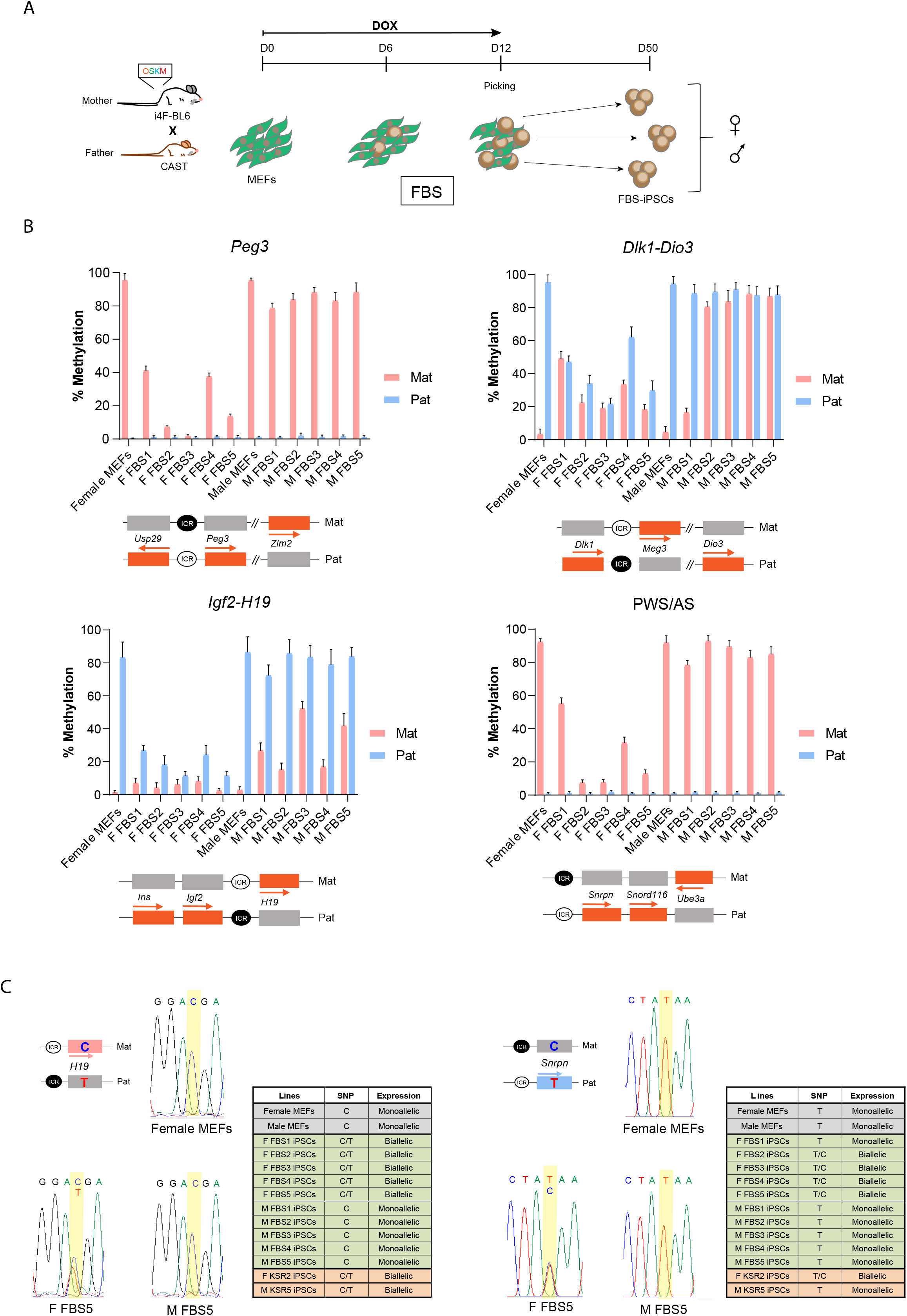
Imprinting methylation defects in F1 FBS-iPSCs. A. Schematic representation of the reprogramming protocol; briefly, a transgenic “reprogrammable” female mouse on a C57BL/6J genetic background (i4F-BL6) was crossed with a *Mus musculus castaneus* (CAST) male mouse to generate E13.5 F1 hybrid embryos from which mouse embryonic fibroblasts (MEFs) were collected. MEFs were reprogrammed by induction of the polycistronic Yamanaka cassette (*Oct4*/*Sox2*/*Klf4*/*c-Myc* - OSKM) in the presence of doxycycline (DOX) for 12 days. Individual mouse induced pluripotent stem cells reprogrammed in Fetal Bovine Serum medium (FBS-iPSCs) clones were picked at day 12 and expanded until approximately day 50. B. Methylation analysis of *Peg3*, *Dlk1-Dio3*, *Igf2-H19* and PWS/AS ICRs in male and female MEFs (note: same data as in Fig. 2A for MEFs), female (F FBS1-5) and male (M FBS1-5) FBS-iPSCs; Each graph represents the mean percentage ± SD methylation levels measured at each CpG within different genomic regions per parental allele for each sample; Scheme on the bottom of each graph represents the normal methylation status of each ICR in the correspondent regions (white circle – unmethylated ICR; black circle – methylated ICR; Mat – maternal allele; Pat – paternal allele; orange rectangles – expressed genes; grey rectangles – silenced genes; regions are not drawn to scale. C. Allelic-specific *H19* and *SNRPN* expression analysis assayed by Sanger sequencing. Chromatograms are shown for female MEFs, F FBS5 and M FBS5 iPSCs; Table summarize the allelic-specific expression for all the FBS-iPSCs as well as female and male MEFs, F KSR2 and M KSR3 iPSCs. Schemes on the left of the female MEFs chromatograms represent the normal imprinting profile of both *H19* and *SNRPN* and the associated SNP for each allele; pink rectangle – maternally *H19* expressed gene; blue rectangle – paternally *Snrpn* expressed gene; grey rectangles – silenced genes; regions are not drawn to scale.

### *Dlk1-Dio3* imprinting is highly labile in iPSCs

We next compared the IMPLICON results obtained for both KSR-and FBS-iPSCs (Fig.5A). This analysis showed that all female iPSCs exhibit a drop in methylation levels at all methylated ICRs irrespective of the medium formulation. Interestingly, two female FBS-iPSC clones (F FBS1 and F FBS4) have consistently milder defects across all the imprinted loci analysed. The unmethylated ICRs in female iPSCs were untouched, but a mild increase at the *Dlk1-Dio3* cluster was consistently seen in a few female FBS-iPSCs (Fig. 5A-B). While male KSR-iPSCs tend to lose methylation at the methylated ICRs, this was found to be less striking than for female KSR/FBS-iPSCs. This data suggests that female sex is the strong driver for hypomethylation defects with the KSR formulation also having an impact. In contrast, male FBS-iPSCs have less deleterious effects on imprinting, except for two clusters with paternal-specific methylated ICRs, *Igf2-H19* and *Dlk1-Dio3*, which gained methylation on the unmethylated ICRs, especially evident for the later locus (Fig. 5A-B).

**Fig. 5.**
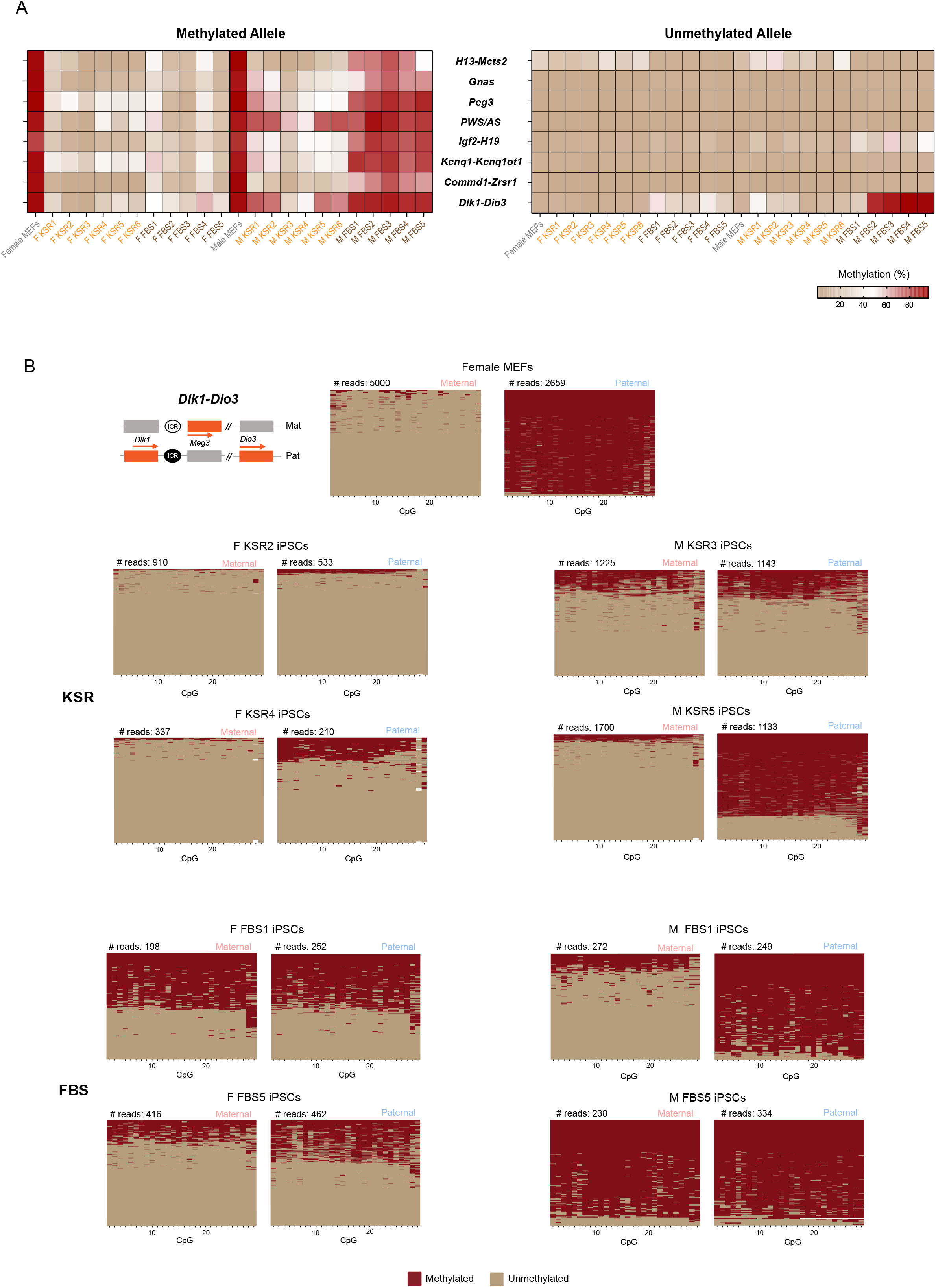
Effect of sex of donor cell and medium conditions on imprinting stability of iPSCs. A. Heatmap representing the percentage of DNA methylation at the methylated (left) and unmethylated (right) ICRs for female and male MEFs as well as female and male KSR-/FBS-iPSCs. ICRs are shown in rows and samples in columns. B. Plots displaying methylated and unmethylated CpGs for each CpG position (in columns) in all the individual reads (in rows) at the *Dlk1-Dio3* ICR in female MEFs, F KSR2, F KSR4, M KSR3, M KSR5, F FBS1, F FBS5, M FBS1, M FBS5 iPSCs; Scheme on the left represents the normal methylation status of the *Dlk1-Dio3* ICR in the correspondent regions (white circle – unmethylated ICR; black circle – methylated ICR; Mat – maternal allele; Pat – paternal allele; orange rectangles –expressed genes; grey rectangles – silenced genes; regions are not drawn to scale).

The *Dlk1-Dio3* cluster was found to be the most labile locus among the 8 imprinted loci investigated in this study. The ICR of the *Dlk1-Dio3* locus showed remarkably diverse methylation outlines ranging from iPSC lines with two unmethylated alleles (F KSR2, F KSR4) to others with two fully methylated alleles (M FBS5), but also including iPSC lines with both alleles partially unmethylated (F FBS1, F FBS5) and a line with correct imprinting status (M FBS1) (Fig. 5A-B; Table S3). We evaluated the impact of these different methylation outcomes in allelic preference and levels of the *Meg3* gene located downstream of the *Dlk1-Dio3* ICR. Biallelic expression was detected in all iPSC lines for which unmethylated reads were found in the two parental alleles. Monoallelic expression was seen for M FBS1 iPSCs and MEFs, which exhibit normal methylation patterns at the *Dlk1-Dio3* ICR. For the hypermethylated M FBS5 this could not be assessed as *Meg3* was not expressed (Fig. S5A). We also tested how the diversity of methylation outlines translate into differences in overall *Meg3* levels measured by RT-qPCR. Although overall *Meg3* levels were consistently higher in KSR versus FBS medium, the levels correlated with the number of unmethylated reads seen for each cell line (Fig. 5B; Fig. S5B). Our data shows that the *Dlk1-Dio3* cluster is highly sensitive to reprogramming conditions and sex of donor cell. This imprinted locus was previously observed to acquire *de novo* methylation in both mouse and human iPSCs (Bar et al., 2017; Stadtfeld et al., 2010), a feature that has been linked to reduced pluripotency of stem cells (Liu et al., 2010; Mo et al., 2015; Stadtfeld et al., 2010). Our results extend the plethora of imprinting defects of this ICR, not only to the maternal unmethylated allele, but also the paternal methylated allele.

### Global DNA methylation differences in iPSCs do not account for their severe imprinting abnormalities

Finally, we evaluated whether global DNA methylation differs between female and male KSR/FBS-iPSCs to understand whether it had an influence on the different imprinting abnormalities found by IMPLICON. First, we monitored the global 5mC levels by liquid chromatography-mass spectrometry (LC-MS) in female and male KSR/FBS-iPSCs and TX 2i ESCs. LC-MS measurements revealed that female iPSCs showed a tendency for lower global levels of 5mC than male iPSCs irrespective of the medium conditions (Fig. 6A). These are consistent with previous findings showing that female mouse ESCs/iPSCs have lower global 5mC levels which is associated with the presence of two active X chromosomes (Milagre et al., 2017; Pasque et al., 2018; Zvetkova et al., 2005). On the other hand, the modest shift towards increased 5mC levels from KSR to FBS conditions for both female and male iPSCs was not found to be statistically significant (Fig. 6A). As expected, both female and male KSR/FBS-iPSCs have considerably higher 5mC levels than the female TX 2i ESCs (Fig. 6A), consistent with minimal methylation levels associated with 2i medium conditions (Leitch et al., 2013; von Meyenn et al., 2016). Therefore, our iPSCs showed relatively mild changes in overall DNA methylation and yet have marked imprinting defects.

**Fig. 6.**
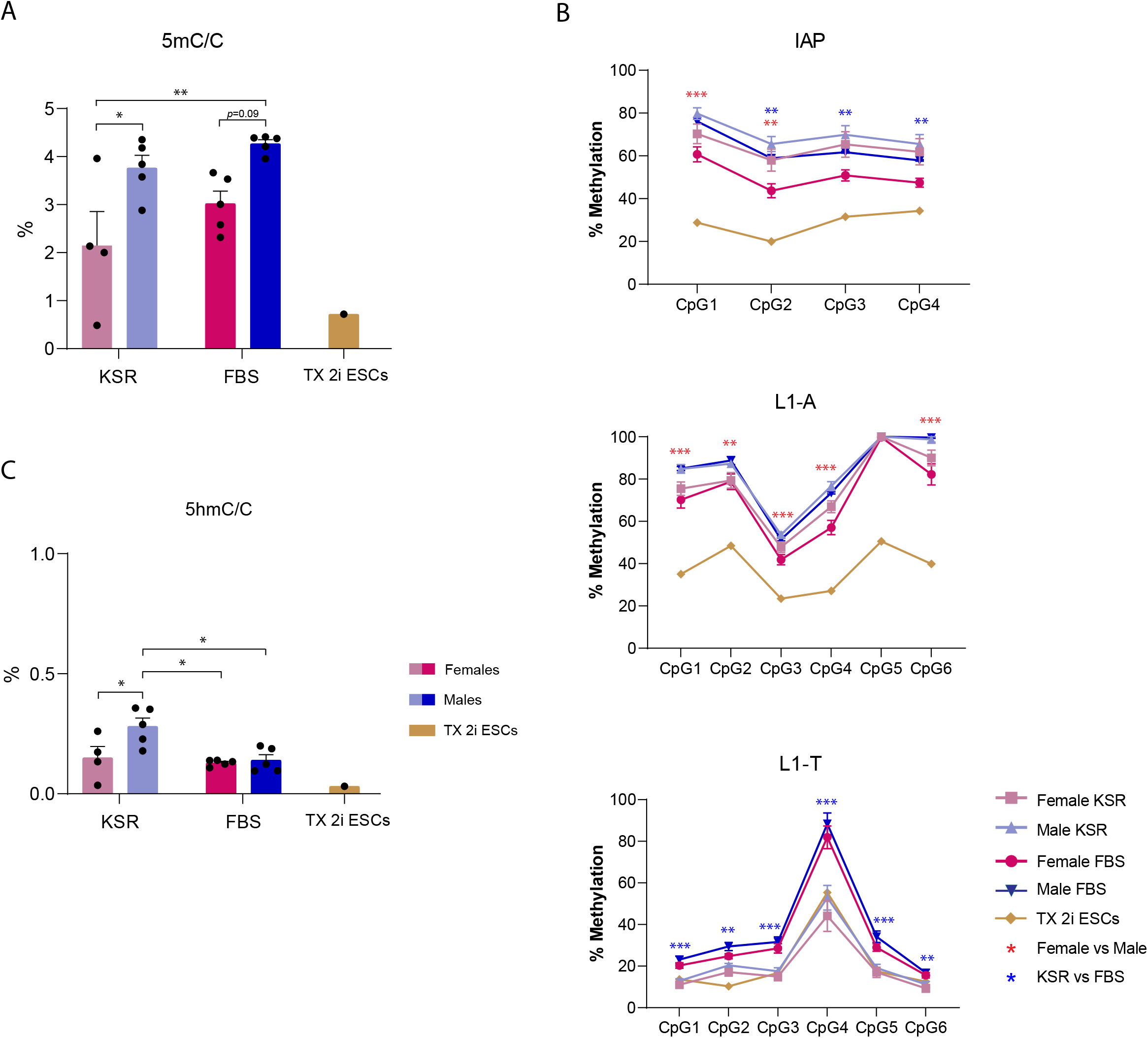
Global methylation changes in female and male KSR-/FBS-derived iPSCs. A. Global 5mC levels measured by LC-MS. Graph represents the average percentage ± SEM of 5mC of the total cytosines in female and male KSR/FBS-iPSCs, as well as in TX 2i ESCs; Individual values is represented by black dots; Statistically significant differences are indicated as * *p* < 0.05; ** *p* < 0.01 (two-way ANOVA followed by Tukey’s multiple comparisons test). B. Percentage CpG methylation measured by bisulfite pyrosequencing for IAP, L1-A and L1-T repetitive elements. Graph represents the average methylation percentages ± SEM at each CpG sampled per repetitive element in female and male KSR/FBS-iPSCs as well as methylation percentages in TX 2i ESCs. Statistically significant differences are indicated as ** *p* < 0.01; *** *p* < 0.001 (two-way ANOVA) with red asterisks meaning statistically significant differences between sex (female vs male) and blue meaning statistically significant differences between medium formulation (KSR vs FBS). C. Global 5hmC levels measured by LC-MS. Graph represents the average percentage ± SEM of 5hmC of the total cytosines in female and male KSR/FBS-iPSCs, as well as in TX 2i ESCs; Individual values is represented by black dots; Statistically significant differences are indicated as * *p* < 0.05 (two-way ANOVA followed by Tukey’s multiple comparisons test).

To complement this analysis, we investigated the methylation levels at abundant retrotransposon elements in the genome, known to be highly methylated in ESCs in serum conditions (Walter et al., 2016). We studied the intracisternal A particle (IAP), the most abundant Long Terminal Repeat (LTR) from the class II of endogenous retroviruses (ERVs) (Stocking and Kozak, 2008) as well as, two subfamilies of young Long Interspersed Nuclear Element 1 (LINE1), LINE1-A (L1-A) and LINE1-T (L1-T), which comprise the most frequent class of non-LTR elements in the mouse genome (Babushok and Kazazian, 2007). By performing quantitative bisulfite-pyrosequencing at these retrotransposon elements (Walter et al., 2016), we showed that each class of retroelements behaved differently according to the sex or medium formulation (Fig. 6B). IAPs suffered a decrease of DNA methylation in female iPSCs, which was potentiated, surprisingly, by FBS medium. L1-A methylation levels were also decreased in female KSR/FBS-iPSCs, (Fig. 6B). These fluctuations in DNA methylation were modest compared to the severe demethylation seen for the TX 2i ESCs (Fig. 6B) as expected (Walter et al., 2016). Interestingly, L1-T levels clearly dropped in both female and male KSR-iPSCs, reaching similarly low methylation levels to TX 2i ESCs (Fig. 6B). In summary, while a modest decrease in the level of 5mC and in methylation of IAPs and L1-A elements could be discerned for female iPSCs (Fig. 6A-B), a strong effect of KSR medium in demethylating L1-T retrotransposons was detected (Fig. 6B). In conclusion, with the exception of L1-T retrotransposons, ICRs are among the most affected loci upon reprogramming with more pronounced effects on DNA methylation than the ones observed at repetitive elements.

To understand the reasons behind global changes in DNA methylation and at imprinted regions in female versus male KSR/FBS-iPSCs, we analysed expression levels of genes involved in DNA methylation and their co-factors (*Dnmt3a*, *Dnmt3b*, *Dnmt3L*, *Dnmt1* and *Uhrf1*), demethylation (*Tet1*, *Tet2* and *Tet3*) and imprinting protection (*Zfp57*, *Trim28* and *Dppa3*). No differences were detected between female and male iPSCs, suggesting that methylation variations at ICRs and globally between sexes are not caused by expression changes in the core DNA methylation/demethylation machinery or imprinting protecting factors (Fig. S6). In contrast, medium formulations had an impact on the expression of these genes: increased levels of *Tet1*, *Tet2* and *Dnmt3a* in KSR medium and *Dnmt3L* and *Uhrf1* in FBS medium (Fig. S6). The increase of *Dnmt3a* levels in KSR medium could be offset by decreased levels of its co-factor *Dnmt3L* (Hata et al., 2002), which could explain why KSR medium does not cause increased DNA methylation. On the other hand, increased levels of the *Tet1* and *Tet2* dioxygenase enzymes in KSR medium could impact by converting 5mC to 5hmC. Interestingly, KSR-iPSCs had higher 5hmC levels than FBS-iPSCs, an effect especially evident for male KSR-iPSCs (Fig. 6C). This effect could be due to the increased levels of *Tet1* and *Tet2*, but also to the presence of ascorbic acid in the KSR formulation which is known to increase the conversion of 5mC to 5hmC by TET dioxygenase enzymes (Blaschke et al., 2013). In conclusion, our analysis suggests that imprinting defects in female iPSCs might be caused by collateral effects due to X chromosome reactivation (Fig. S1D; Fig. S3A) and seems not to rely mainly on changes in the basic DNA methylation machinery or imprinting protection. In contrast, imprinting methylation in male iPSCs might be responsive to medium conditions through fluctuations in the expression and activity of TET dioxygenase enzymes according to medium formulation. Despite the stable protection during development and aging, imprinting is remarkably sensitive to subtle global methylation changes during reprogramming and culture of mouse iPSCs.

## DISCUSSION

In this study, we thoroughly analysed the fidelity of genomic imprinting in mouse iPSCs. By using an established secondary reprogramming system (Abad et al., 2013; Bernardes de Jesus et al., 2018) on murine hybrid cells, we explored the impact of the sex of the donor cell and reprogramming culture conditions on the epigenetic stability of imprinted loci. Our comprehensive analysis defined clear trends associated with the sex of the donor cell and reprogramming conditions: (1) female sex is a strong predictor of hypomethylation imprinting errors in iPSCs; (2) KSR formulation drives hypomethylation defects at many ICRs in iPSCs; (3) classical FBS-based ESC medium conserves correct imprinting in many ICRs in XY cells, but renders paternal methylated ICRs prone to gain methylation on the maternal allele; (4) imprinting is poorly protected from the global DNA methylation fluctuations during reprogramming and expansion of iPSCs. Our systematic analysis provides an explanation for the contrasting results from previous studies (Stadtfeld et al., 2010; Sun et al., 2012; Takikawa et al., 2013; Yagi et al., 2019), when sex of the donor cell (when given) and reprogramming culture conditions are considered (Table S1). This knowledge is important to devise future strategies to create iPSCs devoid of imprinting abnormalities.

The allele-specific IMPLICON creates a final dataset reflecting the original methylated/unmethylated DNA molecules with nucleotide and allelic resolution (Klobucar et al., 2020). This technique allowed us to investigate, for the first time, the heterogeneity of epigenetic states at imprinted regions within iPSC clonal lines. Indeed, we could show that partial hypomethylation defects for certain loci (e.g., *Kcnq1-Kcnq1ot1*, *Gnas*) or particular iPSCs (e.g., male iPSCs) were due to the existence of two major classes of DNA molecules, one with an intact methylated ICR and another with complete erasure of the methylation mark at this element (Fig. S2C; Fig.S4E; Fig. 5B). This suggests that within an iPSC line, a subset of cells retained proper methylation status for a given locus, while the rest lost methylation. Our data points out for intra-heterogeneity of imprinted defects at certain loci within clonal lines which was not previously appreciated.

Female sex is a major predictor of hypomethylation defects at imprinted loci in mouse iPSCs (Fig. 5A). This seems not to be explained by differences in expression levels of genes encoding for the DNA methylation/demethylation enzymes or involved in imprinting protection between female and male iPSCs (Fig. S6) nor an increase in the conversion of 5mC to 5hmC (Fig. 6C). The effect of the female sex might rather be explained by the global wave of DNA demethylation previously reported during reprogramming of female cells associated with X-chromosome reactivation (Milagre et al., 2017; Pasque et al., 2018) from which methylated ICRs do not escape and never recover. During mouse iPSC reprogramming, the inactive X chromosome from somatic cells reactivates to generate iPSCs with two active X chromosomes (Pasque et al., 2014). Higher dosage of X-linked genes may be responsible for this global hypomethylation effect. In particular, the X-linked *Dusp9* gene, which is upregulated in our female iPSCs (Fig. S1D; Table S2) has been implicated in hypomethylation of female stem cells (Choi et al., 2017a). However, other X-linked genes might be involved during reprogramming (Song et al., 2019). While repetitive elements are marginally affected by this global DNA hypomethylation in female iPSCs, our data shows that imprinted loci are highly vulnerable.

Reprogramming under KSR conditions was also a predictor of hypomethylation defects at ICRs in iPSCs. On one hand, imprinted defects occurred even though overall 5mC levels were largely untouched as well as in common repetitive elements (with the exception of L1-T element). On the other hand, hypomethylation at ICRs correlated with elevated *Tet1*/*Tet2* expression levels (Fig. S6) and increase of 5hmC levels globally (Fig. 6C). The effect on 5hmC levels might also be enhanced by the presence of ascorbic acid in the KSR formulation (Stadtfeld et al., 2012) that promotes the activity of TET dioxygenase enzymes and reduces DNA methylation levels (Blaschke et al., 2013). Curiously, ascorbic acid was previously used to successfully recover hypermethylation defects in the *Dlk1-Dio3* cluster in FBS-iPSCs (Stadtfeld et al., 2012). Our results on KSR-iPSCs highlights that the benefits of ascorbic acid in correcting *Dlk1-Dio3* imprinting might be offset by errors built up in other imprinted clusters which should not be underestimated.

The mechanisms behind imprinting defects during reprogramming remain unclear. Recent data suggests the existence of two early global DNA demethylation waves during mouse iPSC reprogramming (Bartoccetti et al., 2020) under KSR conditions with ascorbic acid supplementation. This involved a role for TET1 as an active demethylase leading to the reactivation of germline-specific genes. This resembles germline reprogramming from which imprinting is not protected (Hajkova et al., 2002), which is consistent with KSR-mediated hypomethylation defects at ICRs in an inbred strain observed by Bartoccetti et al. (2020), that we also saw to a slightly greater extent with allele-specific IMPLICON in our hybrid cells. Whether global DNA demethylation and germline-specific gene expression vary with the sex of the cell and medium conditions during reprogramming are important future experiments to elucidate the reasons behind the different outcomes on imprinting fidelity.

Lack of imprinting protection in iPSCs is a clear outcome of our results. Fluctuations in imprinting protecting factors like ZFP57 (a KRAB-zinc finger protein that binds to methylated ICRs), TRIM28 (a co-repressor that links KRAB-ZFP proteins to DNMTs) or DPPA3 (a H3K9me2 binder that reduces TET3 affinity to chromatin and subsequently 5mC conversion to 5hmC) could impair imprinting protection during reprogramming. Previously it was shown that MEFs null for *Zfp57* and *Dppa3* further exacerbate the defects in the resultant iPSCs (McDonald et al., 2016; Xu et al., 2015), highlighting the importance of these protecting factors for imprinting preservation. Particularly crucial might be *Dppa3* since it is not expressed in MEFs (Fig. S6). Indeed, exogenous addition of *Dppa3* had a benefit effect on iPSC reprogramming and *Dlk1-Dio3* imprinting protection (Xu et al., 2015). Addition of *Dppa3* or other protective protein to the Yamanaka cocktail might be an interesting strategy to pursue to avoid imprinting errors in future reprogramming protocols. Alternatively, the use of different stochiometries of the 4 Yamanaka factors or even the exclusion of the *Oct4* from the Yamanaka cocktail might impact positively on imprinting maintenance (Carey et al., 2011; Velychko et al., 2019). This has been suggested for the *Dlk1-Dio3* locus, which has been shown to be particularly susceptible by the variables used in our experimental setup.

Our study clearly points out for recurrent and frequent imprinting errors in iPSCs. Only 1 out of the 22 mouse iPSCs generated (M FBS1 line) had all 8 assessed imprinted regions intact. Apart from a clear negative impact in modelling imprinting disorders, whether defective imprinting could majorly affect the downstream application of iPSCs remains a virgin ground to explore. Our pluripotency assay based on teratoma formation showed that iPSCs with either hypo- or hypermethylation defects (including at the *Dlk1-Dio3* region) could readily generate well-differentiated teratomas. Based on this, we would anticipate that imprinting dysregulation in iPSCs does not impede the major axes of lineage commitment. Imprinting defects might rather affect specific cellular functions or differentiation of defined cellular subtypes, as it has been observed for the role of imprinted gene *IGF2* in hematopoietic commitment (Nishizawa et al., 2016) or *MEG3* in neural differentiation (Mo et al., 2015). These aspects deserve to be attentively studied in the future.

Although neglected as an hallmark of pluripotency (De Los Angeles et al., 2015), the existence of multiple imprinting errors found in cells derived from the first human trial of iPSC-based autologous transplantation (Araki et al., 2019), brings back the topic of imprinting fidelity to the spotlight. Human iPSCs are also known to have frequent imprinting defects (Bar et al., 2017; Ma et al., 2014; Nazor et al., 2012). Our mouse iPSC data might not be directly translated to human iPSCs due to the multiple differences in medium requirements and epigenetic states. For instance, human female iPSCs do not undergo X-chromosome reactivation during reprogramming (Tchieu et al., 2010) and, thus, female sex is not expected to have a major impact on imprinting defects during reprogramming of human iPSCs. Yet, many human female iPSCs/ESCs partially reactivate the inactive X upon prolonged *in vitro* culturing (Mekhoubad et al., 2012; Vallot et al., 2015) and the consequences of that on imprinting are unknown. All in all, our findings in murine iPSCs raise awareness about the importance of testing human iPSCs (or ESCs) for imprinting fidelity using methods such as IMPLICON. Indeed, imprinting screening should be part of a quality control panel to assure safety for the use of human iPSCs for disease modelling approaches, and more importantly, for clinical applications.

## MATERIALS AND METHODS

### Mice strains

The “reprogrammable” transgenic i4-F-BL6 (kind gift from Manuel Serrano, IRB, Barcelona) (Abad et al., 2013), CAST and NSG mice (Shultz et al., 2007) colonies were maintained at the Instituto de Medicina Molecular João Lobo Antunes (iMM JLA) Rodent facility. Animals were housed in a maximum of five per cage in a temperature-controlled room (24°C) with a 14/12hr light/dark cycle. Animals were fed diet *ad libitum*. Animal care and experimental procedures (see MEFs generation and teratoma assay sections) involving mice were carried out in accordance with European Directive 2010/63/EU, transposed to the Portuguese legislation through DL 113/2013. The use of animals has been approved by the Animal Ethics Committee of IMM JLA and by the Portuguese competent authority – Direcção Geral de Alimentação e Veterinária – with license numbers 015229/17 and 023357/19.

### Generation and maintenance of F1 hybrid MEFs

A cross between female i4F-B and male CAST was set up to generate F1 hybrid transgenic E13.5 mouse embryos. MEFs from individual embryos were collected as previously described (Xu, 2005). Genotyping for the inducible Yamanaka cassette and the *rtTA* gene in the *Rosa26* locus was performed using the primers in Table S5. MEFs were cultured in Dulbecco’s modified Eagle’s medium (DMEM; Gibco) containing 10% FBS (Gibco), 2mM GlutaMAX (Gibco) and 100μg/ml penicillin–streptomycin (Gibco). MEFs were expanded and frozen at passage 2 (P2) prior to the reprogramming experiments.

### Reprogramming of MEFs

MEFs at P2 were plated and expanded until reaching confluency of 40% per well in a gelatin-coated 6 well plate and cultured in KSR or FBS iPSC culture conditions (see below) with 1.5 μg/ml DOX (BDClontech). The medium was changed every 48 hours. Individual iPSCs colonies were picked at day 12 after DOX induction using glass cloning cylinders (Sigma-Aldrich) and 0.05% Trypsin-EDTA 1X (Thermo Fisher Scientific) and plated into a previously gelatin-coated 96 well plate on feeders. Each iPSC colony was then transferred subsequently into 24, 12 and 6-well plates and 25 cm^2^ culture flasks. Each iPSC was then continuously passed for 2-4 passages until reaching approximately day 50 after initiation of reprogramming. All cells and reprogramming experiments were performed in an incubator at 37 °C and 5% CO_2_ in normoxia conditions.

### Stem cell culture conditions

KSR-iPSCs were cultured in high-glucose DMEM supplemented with 15% KSR (Gibco), LIF (1000 U/ ml), 1% MEM non-essential amino acids, 0.5% penicillin– streptomycin, 1% glutamax, and 0.2% β-mercaptoethanol. FBS-iPSCs and JM8.F6 ESCs were cultured in standard serum medium conditions which contains high-glucose DMEM supplemented with FBS (15%, Gibco), LIF (1000 U/ ml), 0.5% penicillin– streptomycin, 1% glutamax, and 0.2% β-mercaptoethanol. TX 2i ESCs (Schulz et al., 2014) were cultured in the same medium conditions with the addition of the 2i chemical inhibitors: 3 μM of CHIR99021 and 250 μM of PD0325901 (Sigma).

### Teratoma assay

To evaluate the capacity for teratoma formation, KSR/FBS-iPSCs (F KSR2, F KSR4, M KSR3, M KSR5, F FBS1, M FBS1 and M FBS5) lines were trypsinized and 2 × 10^6^ cells were subcutaneously injected into the flanks of 3-months-old immunocompromised NSG mice. Animals were sacrificed with anesthetic overdose and a necropsy was performed. Subcutaneous tumors were harvested, fixed in 10% neutral-buffered formalin, stained with hematoxylin and eosin and examined by a pathologist blinded to experimental groups using a Leica DM2500 microscope coupled to a Leica MC170 HD microscope camera.

### Immunofluorescence (IF)

Cells previously seeded on gelatin-coated coverslips were fixed with 3% paraformaldehyde for 10 min at room temperature, washed with PBS and permeabilized in 0.5% Triton X-100 in PBS for 4 minutes (min) on ice. A blocking step was performed by incubation with 1 % Bovine Serum Albumin (BSA; Sigma-Aldrich) in PBS for 15 min and subsequently the cells were incubated with primary antibodies for OCT4 (monoclonal clone 7F9.2m Cat# MAB4419 – Millipore, 1:200 dilution), SSEA1 (monoclonal clone MC-480, Cat# MAB4301 – Millipore, 1:100 dilution) and NANOG (polyclonal, Cat# RCAB001P - Cosmo Bio, 1:150 dilution) diluted in 1% BSA/PBS for 45 min. After three washes with PBS, cells were incubated for 45 min with the secondary antibody Cy™3 AffiniPure F(ab’)_2_ Fragment Goat Anti-Mouse IgG (H+L) (Jackson ImmunoResearch Laboratories Inc., 1:200 dilution). DAPI (0.2 mg/ml) was used to stain the DNA and mark the nuclei by incubating at RT for 2 min. Cells were imaged using Zeiss Axio Observer (Carl Zeiss MicroImaging) with 63× oil objective using the filter sets FS43HE and FS49 and digital images were processed using FIJI platform (https://fiji.sc/).

### DNA extraction and bisulfite treatment

Genomic DNA from parental female and male MEFs, respective KSR/FBS-iPSCs and TX 2i ESCs were isolated using conventional phenol:chloroform:isoamyl alcohol extraction. 1μg of genomic DNA was bisulfite converted using the EZ DNA methylation Gold kit (Zymo Research) according to manufacturer’s instructions with column clean-up and eluted in 66μl elution buffer to obtain a final concentration of ~15ng/μl bisulfite converted DNA.

### 5mC/5hmC measurements by liquid chromatography-mass spectrometry

Genomic DNA from both female and male KSR/FBS-iPSCs and TX 2i ESCs was quantified using picogreen assay (Invitrogen), digested using DNA Degradase plus (Zymo Research) overnight at 37°C and analyzed by LC-MS on a LTQ Orbitrap Velos mass spectrometer (Thermo Scientific) fitted with a nanoelectrospray ion-source (Proxeon, Odense, Denmark). Mass spectral data for cytosine, 5-methylcytosine and 5-hydroxymethylcytosine were acquired as previously described (Ficz et al., 2013).

### Bisulfite Pyrosequencing

Bisulfite-converted DNA was amplified using specific primers for L1-A, L1-T and IAP repetitive elements in a 25 μL reaction volume containing 0.4 μM forward and reverse primers, KAPA HiFi HotStart 2X (Roche) and 1μL of bisulfite-treated DNA for KSR/FBS-iPSCs and TX 2i ESCs. PCR conditions and primers are in Table S5. PCR products were purified and annealed with the sequencing primer for pyrosequencing using the PyroMark Q24 (Qiagen).

### IMPLICON library preparation and analysis

IMPLICON was performed as previously described (Klobucar et al., 2020) in parental female and male F1 MEFs, respective KSR/FBS-iPSC lines as well as the TX 2i ESCs. Briefly, following bisulfite conversion, a first PCR amplifies each region per sample in individual reactions, adding adapter sequences, as well as 8 random nucleotides (N_8_) for subsequent data deduplication. PCR conditions and primers for this first step are listed in Table S5. After pooling amplicons for each biological sample and clean-up using AMPure XP magnetic beads (Beckman Coulter), a second PCR completes a sequence-ready library with sample-barcodes for multiplexing. In this PCR reaction, barcoded Illumina adapters are attached to the pooled PCR samples ensuring that each sample pool receives a unique reverse barcoded adapter. Libraries were verified by running 1:30 dilutions on an Agilent bioanalyzer and then sequenced using the Illumina MiSeq platform to generate paired-end 250bp reads using 10% PhIX spike-in as the libraries are of low complexity.

IMPLICON bioinformatics analysis was also performed as described (Klobucar et al., 2020), following the step-by-step guide of data processing analysis in https://github.com/FelixKrueger/IMPLICON. Briefly, data was processed using standard Illumina base-calling pipelines. As the first step in the processing, the first 8 bp of Read 2 were removed and written into the readID of both reads as an in-line barcode, or Unique Molecular Identifier (UMI). This UMI was then later used during the deduplication step with “deduplicate bismark --barcode mapped_file.bam”. Raw sequence reads were then trimmed to remove both poor quality calls and adapters using Trim Galore v0.5.0 (www.bioinformatics.babraham.ac.uk/projects/trim_galore/, Cutadapt version 1.15, parameters: --paired). Trimmed reads were aligned to the mouse reference genome in paired-end mode. Alignments were carried out with Bismark v0.20.0. CpG methylation calls were extracted from the mapping output using the Bismark methylation extractor. Deduplication was then carried out with deduplicate_bismark, using the --barcode option to take UMIs into account (see above). The data was aligned to a hybrid genome of BL6/CAST (the genome was prepared with the SNPsplit package -v0.3.4, https://github.com/FelixKrueger/SNPsplit). Following alignment and deduplication, reads were split allele-specifically with SNPsplit. Aligned read (.bam) files were imported into Seqmonk software (http://www.bioinformatics.babraham.ac.uk/projects/seqmonk) for all downstream analysis. Probes were made for each CpG contained within the amplicon and quantified using the DNA methylation pipeline or total read count options. Downstream analysis was performed using Excel and GraphPad.

From the raw data deposited in GEO under the accession number GSE148067, the reads mapped to the following murine (mm10) genomic coordinates were excluded for consideration in this article for one of the following reasons: (1) regions that fail to reach the coverage threshold for the two parental alleles (> 40 reads); (2) regions not present in all the samples; (3) regions sequenced twice for which only the run with more reads was considered; (4) regions out of the scope of this article: for the samples *NNNN*_4666: Chr1:63261125-63264796, Chr2:174295708-174296349, Chr6:4746303-4746438, Chr6:30736874-30737061, Chr11:22971952-22972131, Chr12:109528329-109528471, Chr15:72809673-72810197, Chr18:36988564-36988740; for the samples *NNNN*_4836: Chr1:63261125-63261262, Chr6:4746303-4746438; (for TX 2i ESCs the region Chr2:174295708-174295902 was also excluded); for the FBS-iPSCs_5069: Chr6:30737609-30737809, Chr7:68144272-68144369, Chr10:13091188-13091317, Chr11:12025411-12025700, Chr17:12742173-12742488, Chr18:12972868-12973155.

### RNA extraction

Total RNA was extracted from cells using NZYOL™ RNA Isolation Reagent (NZYTech) and treated with DNase I (Roche) according to manufacturer’s instructions.

### RT-qPCR

500 ng of DNase-treated RNA was retrotranscribed into cDNA using random hexamers as primers according to the manufacturer’s protocol (Roche Transcriptor High Fidelity cDNA Synthesis Kit). Quantitative real-time PCR was performed using iTaqTM SYBR^®^ Green Supermix (Bio-Rad) in an Applied Biosystems 7500 fast or ViiA 7 equipment to measure expression levels of several genes normalized to the *Gapdh* housekeeping gene. Primers and conditions are listed in Table S5. The relative expression of each RNA of interest was determined using the 2^−ΔΔct^ method.

### RNA-seq

Quality of Dnase I-treated total RNA from biological triplicates of female MEFs, F KSR2, F KSR4, M KSR3, M KSR5 iPSCs and TX 2i ESCs was checked by 2100 Agilent Bioanalyser. Samples with RIN score above 9 were processed. RNA (1μg) was used to generate strand specific polyA 250–300 bp insert cDNA libraries using the bespoke sequencing pipeline at Sanger Institute. Libraries were sequenced with Illumina HiSeq platform using single-end 50bp mode.

RNA-seq raw FastQ data were trimmed with Trim Galore (version 0.6.1, default parameters) and mapped to the mouse GRCm38 genome assembly using Hisat2 version 2.1.0. Differentially expressed genes between female and male KSR-iPSCs were determined using both EdgeR (p-value of 0.05 with multiple testing correction) and intensity difference filter (p-value < 0.05 with multiple testing correction), with the intersection between the two lists giving the high confidence differentially expressed genes. Clustering analysis in Fig. S1B used a Pearson correlation to calculate a distance matrix between all datasets which was then used to construct a neighbour joining tree. Allele-specific alignments were performed by mapping to both CAST_EiJ and C57Bl/6 (GRCm38) genomes, keeping reads that were specific for either genome and excluding those containing conflicting SNP information. Aligned read (bam) files were imported into Seqmonk software (http://www.bioinformatics.babraham.ac.uk/projects/seqmonk) for all downstream analysis using standard parameters. Data was quantified at the mRNA level using strand-specific quantification of mRNA probes using the RNA-seq quantification pipeline in Seqmonk. For the heatmap in Fig. 3A showing allele-specific biases of imprinted genes, selection was based on the following criteria: (1) transcription from a single allele in MEFs (ratio: > 90%:10%), (2) Log2 RPKM > 1 expression in all iPSC replicates; (3) Normalized SNP-specific read counts > 5 in at least two of the three replicates of iPSCs (Table S4).

### Statistics

Statistical analysis used for each experiment is indicated in the respective figure legend with *p*-values indicated or marked as * *p* < 0.05, ** *p* < 0.01 *** *p* < 0.001. The following tests were used: Two-tailed Student *t*-test (Fig. 2B, Fig. S4D and Fig. S5B), two-way ANOVA followed by Tukey’s multiple comparisons test (Fig. 6A-C; Fig. S6).

## ACKNOWLEDGMENTS

We would like to thank Sérgio de Almeida, Miguel Casanova and Inês Milagre for critical reading of the manuscript and the members of the S.T.d.R.’s team for helpful discussions. We also thank Tânia Carvalho and Pedro Ruivo for their help in histological analysis, David Oxley and Judith Webster at Babraham Institute for LC-MS measurements, Bethan Hussey at Sanger Sequencing and Kristina Tabbada at Babraham Institute for assistance with high-throughput sequencing.

Work in S.T.d.R.’s team was supported by Fundação para a Ciência e Tecnologia (FCT) Ministério da Ciência, Tecnologia e Ensino Superior (MCTES), Portugal [PTDC/BEX-BCM/2612/2014, PTDC/BIA-MOL/29320/2017 IC&DT]; S.T.d.R. has a CEECUIND/01234/207 assistant research contract and A.C.R. has a PhD SFRH/BD/137099/2018 fellowship from FCT/MCTES. MAE-M is supported by a BBSRC Discovery Fellowship (BB/T009713/1).

## REFERENCES

Abad, M., Mosteiro, L., Pantoja, C., Canamero, M., Rayon, T., Ors, I., Grana, O., Megias, D., Dominguez, O., Martinez, D., et al. (2013). Reprogramming in vivo produces teratomas and iPS cells with totipotency features. Nature 502, 340–345.

Araki, H., Miura, F., Watanabe, A., Morinaga, C., Kitaoka, F., Kitano, Y., Sakai, N., Shibata, Y., Terada, M., Goto, S., et al. (2019). Base-Resolution Methylome of Retinal Pigment Epithelial Cells Used in the First Trial of Human Induced Pluripotent Stem Cell-Based Autologous Transplantation. Stem Cell Reports 13, 761–774.

Avior, Y., Eggan, K., and Benvenisty, N. (2019). Cancer-Related Mutations Identified in Primed and Naive Human Pluripotent Stem Cells. Cell Stem Cell 25, 456–461.

Babushok, D.V., and Kazazian, H.H., Jr. (2007). Progress in understanding the biology of the human mutagen LINE-1. Hum Mutat 28, 527–539.

Bar-Nur, O., Russ, H.A., Efrat, S., and Benvenisty, N. (2011). Epigenetic memory and preferential lineage-specific differentiation in induced pluripotent stem cells derived from human pancreatic islet beta cells. Cell Stem Cell 9, 17–23.

Bar, S., Schachter, M., Eldar-Geva, T., and Benvenisty, N. (2017). Large-Scale Analysis of Loss of Imprinting in Human Pluripotent Stem Cells. Cell Rep 19, 957–968.

Bartoccetti, M., van der Veer, B.K., Luo, X., Khoueiry, R., She, P., Bajaj, M., Xu, J., Janiszewski, A., Thienpont, B., Pasque, V., et al. (2020). Regulatory Dynamics of Tet1 and Oct4 Resolve Stages of Global DNA Demethylation and Transcriptomic Changes in Reprogramming. Cell Rep 30, 3948.

Ben-David, U., Arad, G., Weissbein, U., Mandefro, B., Maimon, A., Golan-Lev, T., Narwani, K., Clark, A.T., Andrews, P.W., Benvenisty, N., et al. (2014). Aneuploidy induces profound changes in gene expression, proliferation and tumorigenicity of human pluripotent stem cells. Nat Commun 5, 4825.

Bernardes de Jesus, B., Marinho, S.P., Barros, S., Sousa-Franco, A., Alves-Vale, C., Carvalho, T., and Carmo-Fonseca, M. (2018). Silencing of the lncRNA Zeb2-NAT facilitates reprogramming of aged fibroblasts and safeguards stem cell pluripotency. Nat Commun 9, 94.

Blaschke, K., Ebata, K.T., Karimi, M.M., Zepeda-Martinez, J.A., Goyal, P., Mahapatra, S., Tam, A., Laird, D.J., Hirst, M., Rao, A., et al. (2013). Vitamin C induces Tet-dependent DNA demethylation and a blastocyst-like state in ES cells. Nature 500, 222–226.

Bock, C., Kiskinis, E., Verstappen, G., Gu, H., Boulting, G., Smith, Z.D., Ziller, M., Croft, G.F., Amoroso, M.W., Oakley, D.H., et al. (2011). Reference Maps of human ES and iPS cell variation enable high-throughput characterization of pluripotent cell lines. Cell 144, 439–452.

Buganim, Y., Faddah, D.A., and Jaenisch, R. (2013). Mechanisms and models of somatic cell reprogramming. Nat Rev Genet 14, 427–439.

Carey, B.W., Markoulaki, S., Hanna, J.H., Faddah, D.A., Buganim, Y., Kim, J., Ganz, K., Steine, E.J., Cassady, J.P., Creyghton, M.P., et al. (2011). Reprogramming factor stoichiometry influences the epigenetic state and biological properties of induced pluripotent stem cells. Cell Stem Cell 9, 588–598.

Cassidy, S.B., Schwartz, S., Miller, J.L., and Driscoll, D.J. (2012). Prader-Willi syndrome. Genet Med 14, 10–26.

Choi, J., Clement, K., Huebner, A.J., Webster, J., Rose, C.M., Brumbaugh, J., Walsh, R.M., Lee, S., Savol, A., Etchegaray, J.P., et al. (2017a). DUSP9 Modulates DNA Hypomethylation in Female Mouse Pluripotent Stem Cells. Cell Stem Cell 20, 706–719 e707.

Choi, J., Huebner, A.J., Clement, K., Walsh, R.M., Savol, A., Lin, K., Gu, H., Di Stefano, B., Brumbaugh, J., Kim, S.Y., et al. (2017b). Prolonged Mek1/2 suppression impairs the developmental potential of embryonic stem cells. Nature 548, 219–223.

De Los Angeles, A., Ferrari, F., Xi, R., Fujiwara, Y., Benvenisty, N., Deng, H., Hochedlinger, K., Jaenisch, R., Lee, S., Leitch, H.G., et al. (2015). Hallmarks of pluripotency. Nature 525, 469–478.

Ficz, G., Hore, T.A., Santos, F., Lee, H.J., Dean, W., Arand, J., Krueger, F., Oxley, D., Paul, Y.L., Walter, J., et al. (2013). FGF signaling inhibition in ESCs drives rapid genome-wide demethylation to the epigenetic ground state of pluripotency. Cell Stem Cell 13, 351–359.

Gore, A., Li, Z., Fung, H.L., Young, J.E., Agarwal, S., Antosiewicz-Bourget, J., Canto, I., Giorgetti, A., Israel, M.A., Kiskinis, E., et al. (2011). Somatic coding mutations in human induced pluripotent stem cells. Nature 471, 63–67.

Habibi, E., Brinkman, A.B., Arand, J., Kroeze, L.I., Kerstens, H.H., Matarese, F., Lepikhov, K., Gut, M., Brun-Heath, I., Hubner, N.C., et al. (2013). Whole-genome bisulfite sequencing of two distinct interconvertible DNA methylomes of mouse embryonic stem cells. Cell Stem Cell 13, 360–369.

Hajkova, P., Erhardt, S., Lane, N., Haaf, T., El-Maarri, O., Reik, W., Walter, J., and Surani, M.A. (2002). Epigenetic reprogramming in mouse primordial germ cells. Mech Dev 117, 15–23.

Hata, K., Okano, M., Lei, H., and Li, E. (2002). Dnmt3L cooperates with the Dnmt3 family of de novo DNA methyltransferases to establish maternal imprints in mice. Development 129, 1983–1993.

Huang, K., Shen, Y., Xue, Z., Bibikova, M., April, C., Liu, Z., Cheng, L., Nagy, A., Pellegrini, M., Fan, J.B., et al. (2014). A panel of CpG methylation sites distinguishes human embryonic stem cells and induced pluripotent stem cells. Stem Cell Reports 2, 36–43.

Kajiwara, M., Aoi, T., Okita, K., Takahashi, R., Inoue, H., Takayama, N., Endo, H., Eto, K., Toguchida, J., Uemoto, S., et al. (2012). Donor-dependent variations in hepatic differentiation from human-induced pluripotent stem cells. Proc Natl Acad Sci U S A 109, 12538–12543.

Kim, K., Zhao, R., Doi, A., Ng, K., Unternaehrer, J., Cahan, P., Huo, H., Loh, Y.H., Aryee, M.J., Lensch, M.W., et al. (2011). Donor cell type can influence the epigenome and differentiation potential of human induced pluripotent stem cells. Nat Biotechnol 29, 1117–1119.

Klobucar, T., Kreibich, E., Krueger, F., Arez, M., Polvora-Brandao, D., von Meyenn, F., da Rocha, S.T., and Eckersley-Maslin, M. (2020). IMPLICON: an ultra-deep sequencing method to uncover DNA methylation at imprinted regions. Nucleic Acids Res 48, e92.

Koyanagi-Aoi, M., Ohnuki, M., Takahashi, K., Okita, K., Noma, H., Sawamura, Y., Teramoto, I., Narita, M., Sato, Y., Ichisaka, T., et al. (2013). Differentiation-defective phenotypes revealed by large-scale analyses of human pluripotent stem cells. Proc Natl Acad Sci U S A 110, 20569–20574.

Lee, J., Matsuzawa, A., Shiura, H., Sutani, A., and Ishino, F. (2018). Preferable in vitro condition for maintaining faithful DNA methylation imprinting in mouse embryonic stem cells. Genes Cells 23, 146–160.

Leitch, H.G., McEwen, K.R., Turp, A., Encheva, V., Carroll, T., Grabole, N., Mansfield, W., Nashun, B., Knezovich, J.G., Smith, A., et al. (2013). Naive pluripotency is associated with global DNA hypomethylation. Nat Struct Mol Biol 20, 311–316.

Lister, R., Pelizzola, M., Kida, Y.S., Hawkins, R.D., Nery, J.R., Hon, G., Antosiewicz-Bourget, J., O’Malley, R., Castanon, R., Klugman, S., et al. (2011). Hotspots of aberrant epigenomic reprogramming in human induced pluripotent stem cells. Nature 471, 68–73.

Liu, L., Luo, G.Z., Yang, W., Zhao, X., Zheng, Q., Lv, Z., Li, W., Wu, H.J., Wang, L., Wang, X.J., et al. (2010). Activation of the imprinted Dlk1-Dio3 region correlates with pluripotency levels of mouse stem cells. J Biol Chem 285, 19483–19490.

Ma, H., Morey, R., O’Neil, R.C., He, Y., Daughtry, B., Schultz, M.D., Hariharan, M., Nery, J.R., Castanon, R., Sabatini, K., et al. (2014). Abnormalities in human pluripotent cells due to reprogramming mechanisms. Nature 511, 177–183.

Mandai, M., Fujii, M., Hashiguchi, T., Sunagawa, G.A., Ito, S.I., Sun, J., Kaneko, J., Sho, J., Yamada, C., and Takahashi, M. (2017). iPSC-Derived Retina Transplants Improve Vision in rd1 End-Stage Retinal-Degeneration Mice. Stem Cell Reports 8, 1112–1113.

Maranga, C., Fernandes, T.G., Bekman, E., and da Rocha, S.T. (2020). Angelman syndrome: a journey through the brain. FEBS J 287, 2154–2175.

McDonald, C.M., Liu, L., Xiao, L., Schaniel, C., and Li, X. (2016). Genomic imprinting defect in Zfp57 mutant iPS cell lines. Stem Cell Res 16, 259–263.

Mekhoubad, S., Bock, C., de Boer, A.S., Kiskinis, E., Meissner, A., and Eggan, K. (2012). Erosion of dosage compensation impacts human iPSC disease modeling. Cell Stem Cell 10, 595–609.

Milagre, I., Stubbs, T.M., King, M.R., Spindel, J., Santos, F., Krueger, F., Bachman, M., Segonds-Pichon, A., Balasubramanian, S., Andrews, S.R., et al. (2017). Gender Differences in Global but Not Targeted Demethylation in iPSC Reprogramming. Cell Rep 18, 1079–1089.

Mo, C.F., Wu, F.C., Tai, K.Y., Chang, W.C., Chang, K.W., Kuo, H.C., Ho, H.N., Chen, H.F., and Lin, S.P. (2015). Loss of non-coding RNA expression from the DLK1-DIO3 imprinted locus correlates with reduced neural differentiation potential in human embryonic stem cell lines. Stem Cell Res Ther 6, 1.

Nazor, K.L., Altun, G., Lynch, C., Tran, H., Harness, J.V., Slavin, I., Garitaonandia, I., Muller, F.J., Wang, Y.C., Boscolo, F.S., et al. (2012). Recurrent variations in DNA methylation in human pluripotent stem cells and their differentiated derivatives. Cell Stem Cell 10, 620–634.

Nishizawa, M., Chonabayashi, K., Nomura, M., Tanaka, A., Nakamura, M., Inagaki, A., Nishikawa, M., Takei, I., Oishi, A., Tanabe, K., et al. (2016). Epigenetic Variation between Human Induced Pluripotent Stem Cell Lines Is an Indicator of Differentiation Capacity. Cell Stem Cell 19, 341–354.

Ohi, Y., Qin, H., Hong, C., Blouin, L., Polo, J.M., Guo, T., Qi, Z., Downey, S.L., Manos, P.D., Rossi, D.J., et al. (2011). Incomplete DNA methylation underlies a transcriptional memory of somatic cells in human iPS cells. Nat Cell Biol 13, 541–549.

Pasque, V., Karnik, R., Chronis, C., Petrella, P., Langerman, J., Bonora, G., Song, J., Vanheer, L., Sadhu Dimashkie, A., Meissner, A., et al. (2018). X Chromosome Dosage Influences DNA Methylation Dynamics during Reprogramming to Mouse iPSCs. Stem Cell Reports 10, 1537–1550.

Pasque, V., Tchieu, J., Karnik, R., Uyeda, M., Sadhu Dimashkie, A., Case, D., Papp, B., Bonora, G., Patel, S., Ho, R., et al. (2014). X chromosome reactivation dynamics reveal stages of reprogramming to pluripotency. Cell 159, 1681–1697.

Polo, J.M., Liu, S., Figueroa, M.E., Kulalert, W., Eminli, S., Tan, K.Y., Apostolou, E., Stadtfeld, M., Li, Y., Shioda, T., et al. (2010). Cell type of origin influences the molecular and functional properties of mouse induced pluripotent stem cells. Nat Biotechnol 28, 848–855.

Ruiz, S., Diep, D., Gore, A., Panopoulos, A.D., Montserrat, N., Plongthongkum, N., Kumar, S., Fung, H.L., Giorgetti, A., Bilic, J., et al. (2012). Identification of a specific reprogramming-associated epigenetic signature in human induced pluripotent stem cells. Proc Natl Acad Sci U S A 109, 16196–16201.

Schiebinger, G., Shu, J., Tabaka, M., Cleary, B., Subramanian, V., Solomon, A., Gould, J., Liu, S., Lin, S., Berube, P., et al. (2019). Optimal-Transport Analysis of Single-Cell Gene Expression Identifies Developmental Trajectories in Reprogramming. Cell 176, 1517.

Schulz, E.G., Meisig, J., Nakamura, T., Okamoto, I., Sieber, A., Picard, C., Borensztein, M., Saitou, M., Bluthgen, N., and Heard, E. (2014). The two active X chromosomes in female ESCs block exit from the pluripotent state by modulating the ESC signaling network. Cell Stem Cell 14, 203–216.

Shi, Y., Inoue, H., Wu, J.C., and Yamanaka, S. (2017). Induced pluripotent stem cell technology: a decade of progress. Nat Rev Drug Discov 16, 115–130.

Shultz, L.D., Ishikawa, F., and Greiner, D.L. (2007). Humanized mice in translational biomedical research. Nat Rev Immunol 7, 118–130.

Soellner, L., Begemann, M., Mackay, D.J., Gronskov, K., Tumer, Z., Maher, E.R., Temple, I.K., Monk, D., Riccio, A., Linglart, A., et al. (2017). Recent Advances in Imprinting Disorders. Clin Genet 91, 3–13.

Song, J., Janiszewski, A., De Geest, N., Vanheer, L., Talon, I., El Bakkali, M., Oh, T., and Pasque, V. (2019). X-Chromosome Dosage Modulates Multiple Molecular and Cellular Properties of Mouse Pluripotent Stem Cells Independently of Global DNA Methylation Levels. Stem Cell Reports 12, 333–350.

Stadtfeld, M., Apostolou, E., Akutsu, H., Fukuda, A., Follett, P., Natesan, S., Kono, T., Shioda, T., and Hochedlinger, K. (2010). Aberrant silencing of imprinted genes on chromosome 12qF1 in mouse induced pluripotent stem cells. Nature 465, 175–181.

Stadtfeld, M., Apostolou, E., Ferrari, F., Choi, J., Walsh, R.M., Chen, T., Ooi, S.S., Kim, S.Y., Bestor, T.H., Shioda, T., et al. (2012). Ascorbic acid prevents loss of Dlk1-Dio3 imprinting and facilitates generation of all-iPS cell mice from terminally differentiated B cells. Nature genetics 44, 398–405, S391–392.

Stocking, C., and Kozak, C.A. (2008). Murine endogenous retroviruses. Cell Mol Life Sci 65, 3383–3398.

Strogantsev, R., Krueger, F., Yamazawa, K., Shi, H., Gould, P., Goldman-Roberts, M., McEwen, K., Sun, B., Pedersen, R., and Ferguson-Smith, A.C. (2015). Allele-specific binding of ZFP57 in the epigenetic regulation of imprinted and non-imprinted monoallelic expression. Genome Biol 16, 112.

Sun, B., Ito, M., Mendjan, S., Ito, Y., Brons, I.G., Murrell, A., Vallier, L., Ferguson-Smith, A.C., and Pedersen, R.A. (2012). Status of genomic imprinting in epigenetically distinct pluripotent stem cells. Stem Cells 30, 161–168.

Takahashi, K., and Yamanaka, S. (2006). Induction of pluripotent stem cells from mouse embryonic and adult fibroblast cultures by defined factors. Cell 126, 663–676.

Takikawa, S., Ray, C., Wang, X., Shamis, Y., Wu, T.Y., and Li, X. (2013). Genomic imprinting is variably lost during reprogramming of mouse iPS cells. Stem Cell Res 11, 861–873.

Tchieu, J., Kuoy, E., Chin, M.H., Trinh, H., Patterson, M., Sherman, S.P., Aimiuwu, O., Lindgren, A., Hakimian, S., Zack, J.A., et al. (2010). Female human iPSCs retain an inactive X chromosome. Cell Stem Cell 7, 329–342.

Tonge, P.D., Corso, A.J., Monetti, C., Hussein, S.M., Puri, M.C., Michael, I.P., Li, M., Lee, D.S., Mar, J.C., Cloonan, N., et al. (2014). Divergent reprogramming routes lead to alternative stem-cell states. Nature 516, 192–197.

Tucci, V., Isles, A.R., Kelsey, G., Ferguson-Smith, A.C., and Erice Imprinting, G. (2019). Genomic Imprinting and Physiological Processes in Mammals. Cell 176, 952–965.

Vallot, C., Ouimette, J.F., Makhlouf, M., Feraud, O., Pontis, J., Come, J., Martinat, C., Bennaceur-Griscelli, A., Lalande, M., and Rougeulle, C. (2015). Erosion of X Chromosome Inactivation in Human Pluripotent Cells Initiates with XACT Coating and Depends on a Specific Heterochromatin Landscape. Cell Stem Cell 16, 533–546.

Velychko, S., Adachi, K., Kim, K.P., Hou, Y., MacCarthy, C.M., Wu, G., and Scholer, H.R. (2019). Excluding Oct4 from Yamanaka Cocktail Unleashes the Developmental Potential of iPSCs. Cell Stem Cell 25, 737–753 e734.

von Meyenn, F., Iurlaro, M., Habibi, E., Liu, N.Q., Salehzadeh-Yazdi, A., Santos, F., Petrini, E., Milagre, I., Yu, M., Xie, Z., et al. (2016). Impairment of DNA Methylation Maintenance Is the Main Cause of Global Demethylation in Naive Embryonic Stem Cells. Molecular cell 62, 983.

Walter, M., Teissandier, A., Perez-Palacios, R., and Bourc’his, D. (2016). An epigenetic switch ensures transposon repression upon dynamic loss of DNA methylation in embryonic stem cells. Elife 5.

Wang, K.H., Kupa, J., Duffy, K.A., and Kalish, J.M. (2019). Diagnosis and Management of Beckwith-Wiedemann Syndrome. Front Pediatr 7, 562.

Xie, W., Barr, C.L., Kim, A., Yue, F., Lee, A.Y., Eubanks, J., Dempster, E.L., and Ren, B. (2012). Base-resolution analyses of sequence and parent-of-origin dependent DNA methylation in the mouse genome. Cell 148, 816–831.

Xu, J. (2005). Preparation, culture, and immortalization of mouse embryonic fibroblasts. Curr Protoc Mol Biol Chapter 28, Unit 28 21.

Xu, X., Smorag, L., Nakamura, T., Kimura, T., Dressel, R., Fitzner, A., Tan, X., Linke, M., Zechner, U., Engel, W., et al. (2015). Dppa3 expression is critical for generation of fully reprogrammed iPS cells and maintenance of Dlk1-Dio3 imprinting. Nat Commun 6, 6008.

Yagi, M., Kabata, M., Ukai, T., Ohta, S., Tanaka, A., Shimada, Y., Sugimoto, M., Araki, K., Okita, K., Woltjen, K., et al. (2019). De Novo DNA Methylation at Imprinted Loci during Reprogramming into Naive and Primed Pluripotency. Stem Cell Reports 12, 1113–1128.

Yamanaka, S. (2009). Elite and stochastic models for induced pluripotent stem cell generation. Nature 460, 49–52.

Zvetkova, I., Apedaile, A., Ramsahoye, B., Mermoud, J.E., Crompton, L.A., John, R., Feil, R., and Brockdorff, N. (2005). Global hypomethylation of the genome in XX embryonic stem cells. Nature genetics 37, 1274–1279.

