## Supplementary Information for "Sex of donor cell and reprogramming conditions predict the extent and nature of imprinting defects in mouse iPSCs"

### Figure S1

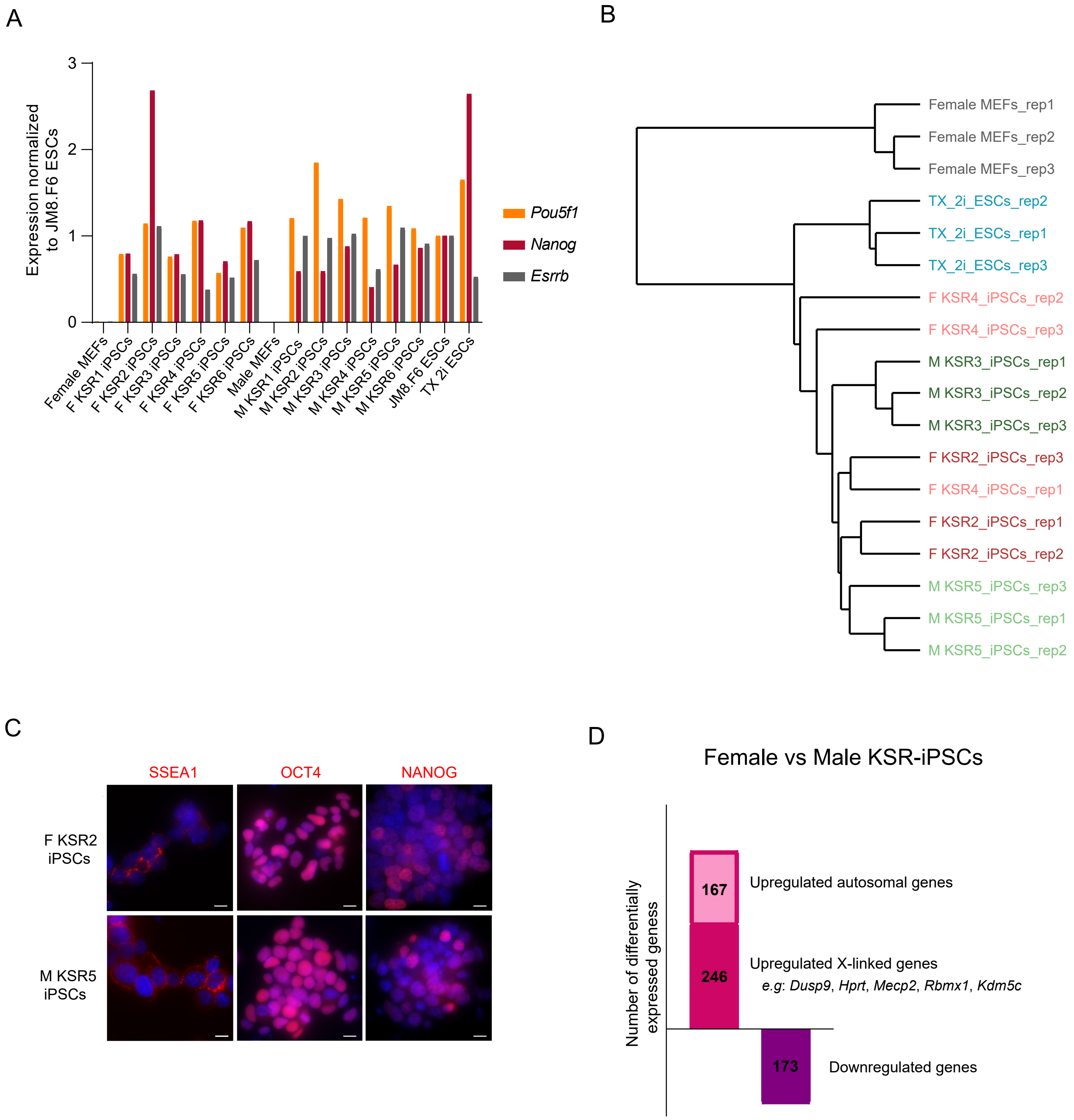
A
3
2
1
0
Expression normalized
to JM8.F6 ESCs
Pou5f1
Nanog
Esrrb
Female MEFs
F KSR1 iPSCs
F KSR2 iPSCs
F KSR3 iPSCs
F KSR4 iPSCs
F KSR5 iPSCs
F KSR6 iPSCs
Male MEFs
M KSR1 iPSCs
M KSR2 iPSCs
M KSR3 iPSCs
M KSR4 iPSCs
M KSR5 iPSCs
M KSR6 iPSCs
JM8.F6 ESCs
TX 2i ESCs
B
Female MEFs_rep1
Female MEFs_rep2
Female MEFs_rep3
TX_2i_ESCs_rep2
TX_2i_ESCs_rep1
TX_2i_ESCs_rep3
F KSR4_iPSCs_rep2
F KSR4_iPSCs_rep3
M KSR3_iPSCs_rep1
M KSR3_iPSCs_rep2
M KSR3_iPSCs_rep3
F KSR2_iPSCs_rep3
F KSR4_iPSCs_rep1
F KSR2_iPSCs_rep1
F KSR2_iPSCs_rep2
M KSR5_iPSCs_rep3
M KSR5_iPSCs_rep1
M KSR5_iPSCs_rep2
C
SSEA1
OCT4
NANOG
F KSR2
iPSCs
M KSR5
iPSCs
D
Female vs Male KSR-iPSCs
Number of differentially
expressed geness
167
Upregulated autosomal genes
246
Upregulated X-linked genes
e.g: Dusp9, Hprt, Mecp2, Rbmx1, Kdm5c
173
Downregulated genes

### Figure S2

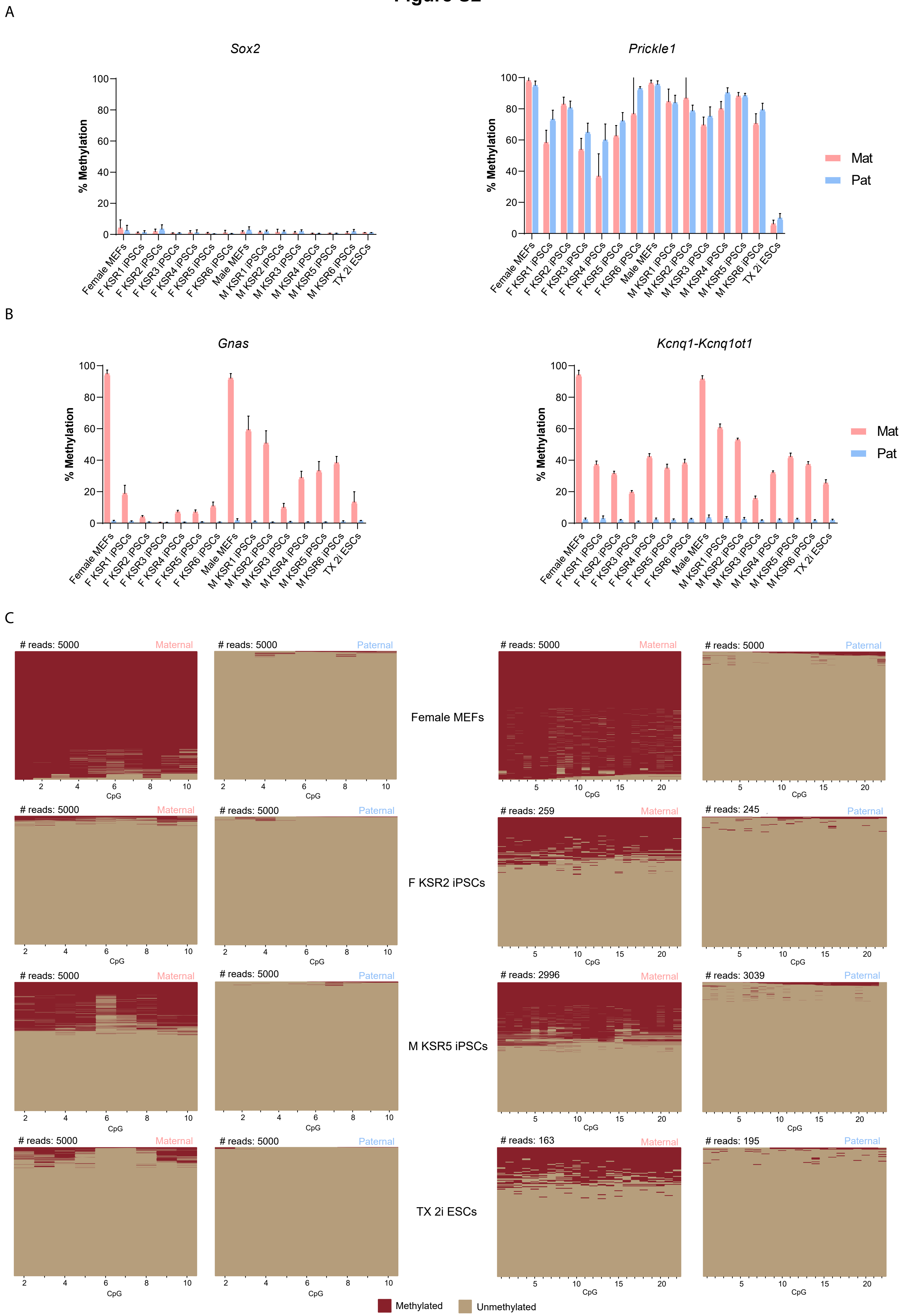
A
Sox2
Prickle1
% Methylation
Mat
Pat
Female MEFs
F KSR1 iPSCs
F KSR2 iPSCs
F KSR3 iPSCs
F KSR4 iPSCs
F KSR5 iPSCs
F KSR6 iPSCs
Male MEFs
M KSR1 iPSCs
M KSR2 iPSCs
M KSR3 iPSCs
M KSR4 iPSCs
M KSR5 iPSCs
M KSR6 iPSCs
TX 2i ESCs
B
Gnas
Kcnq1-Kcnq1ot1
C
### reads: 5000
### reads: 259
### reads: 245
### reads: 2996
### reads: 3039
### reads: 163
### reads: 195
Maternal
Paternal
CpG
Female MEFs
F KSR2 iPSCs
M KSR5 iPSCs
TX 2i ESCs
Methylated
Unmethylated

### Figure S3

A

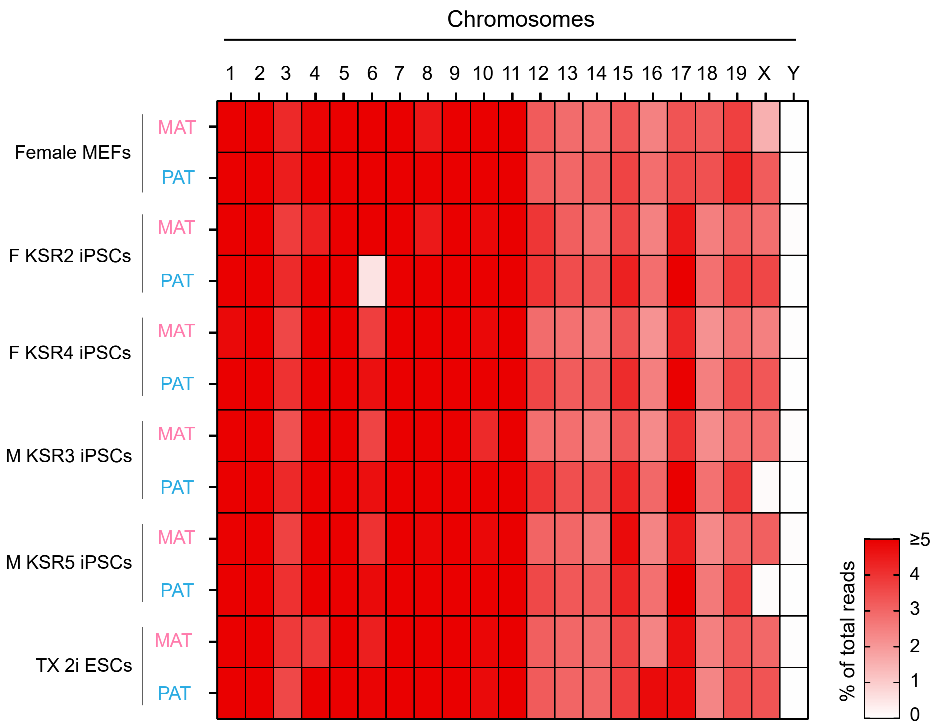
Chromosomes
1 2 3 4 5 6 7 8 9 10 11 12 13 14 15 16 17 18 19 X Y
Female MEFs
MAT
PAT
F KSR2 iPSCs
MAT
PAT
F KSR4 iPSCs
MAT
PAT
M KSR3 iPSCs
MAT
PAT
M KSR5 iPSCs
MAT
PAT
TX 2i ESCs
MAT
PAT
% of total reads
≥5
4
3
2
1
0

B

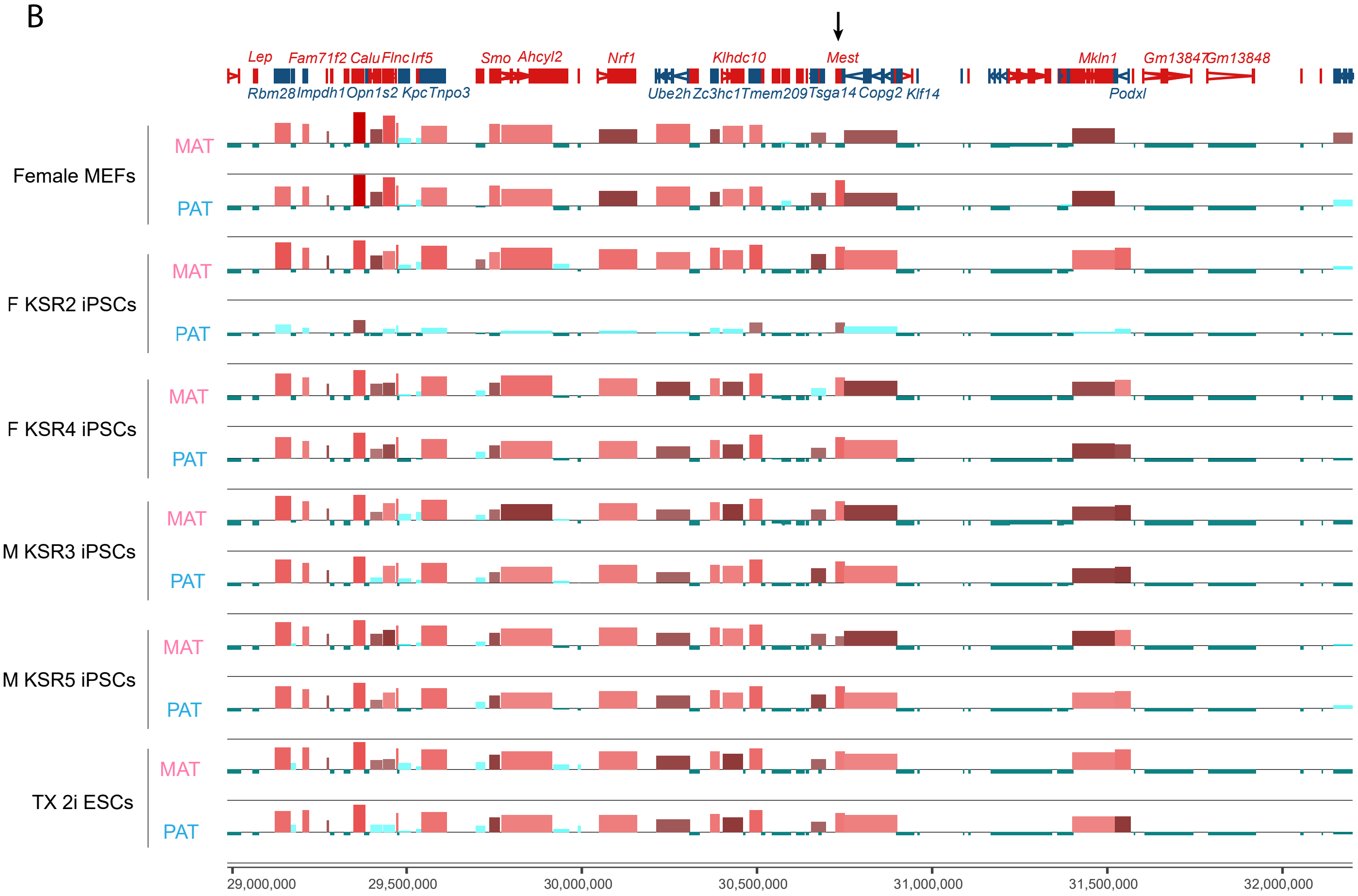
Lep Fam71f2 Calu Flnc Irf5 Smo Ahcyl2 Nrf1 Klhdc10 Mest Mkln1 Gm13847 Gm13848
Rbm28 Impdh1 Opn1s2 Kpc Tnpo3 Ube2h Zc3hc1 Tmem209 Tsga14 Copg2 Klf14 Podxl
Female MEFs
MAT
PAT
F KSR2 iPSCs
MAT
PAT
F KSR4 iPSCs
MAT
PAT
M KSR3 iPSCs
MAT
PAT
M KSR5 iPSCs
MAT
PAT
TX 2i ESCs
MAT
PAT
29,000,000
29,500,000
30,000,000
30,500,000
31,000,000
31,500,000
32,000,000

### Figure S4

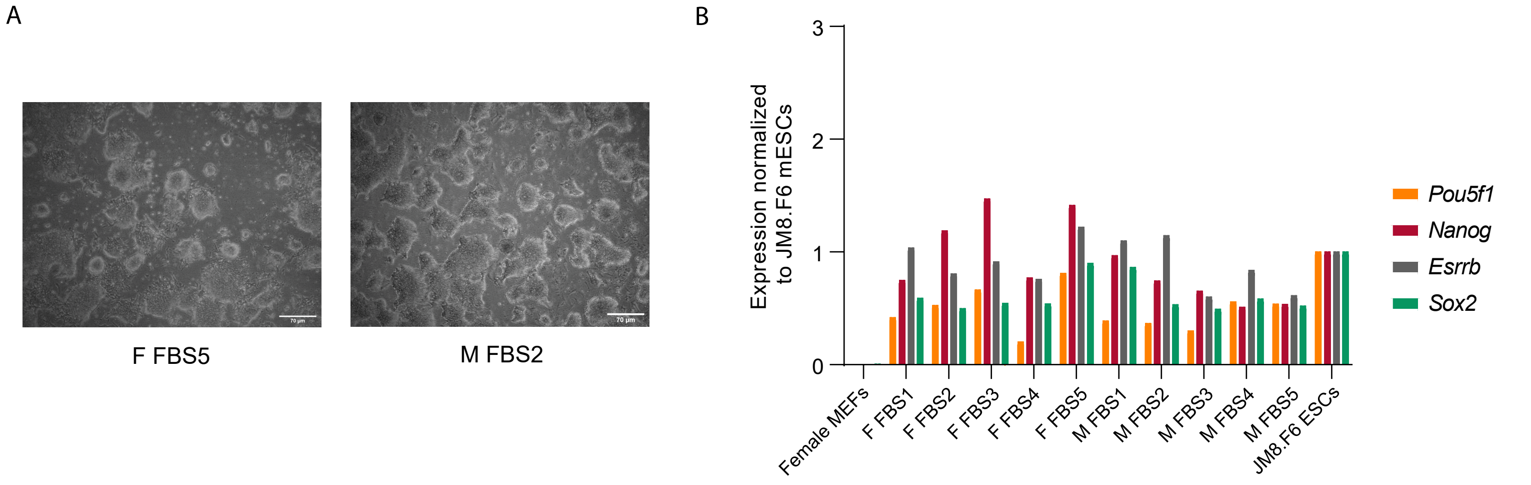


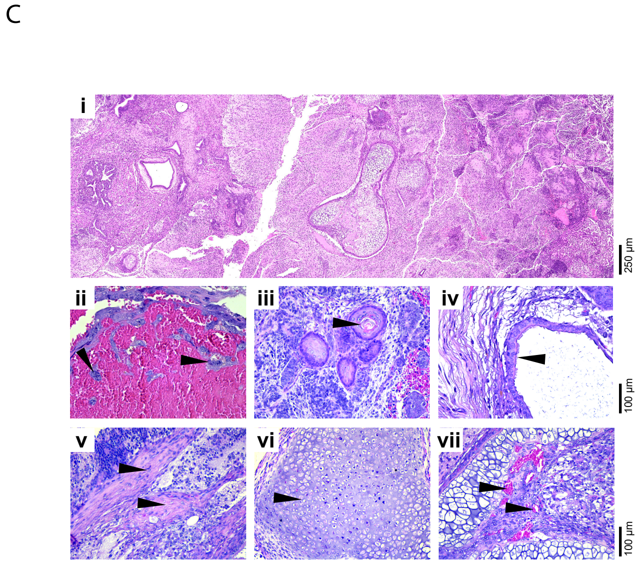


| iPSCs lines | Ectoderm | Mesoderm | Endoderm |
|---|---|---|---|
| F FBS1 | + | + | + |
| M FBS1 | + | + | + |
| M FBS5 | + | + | + |

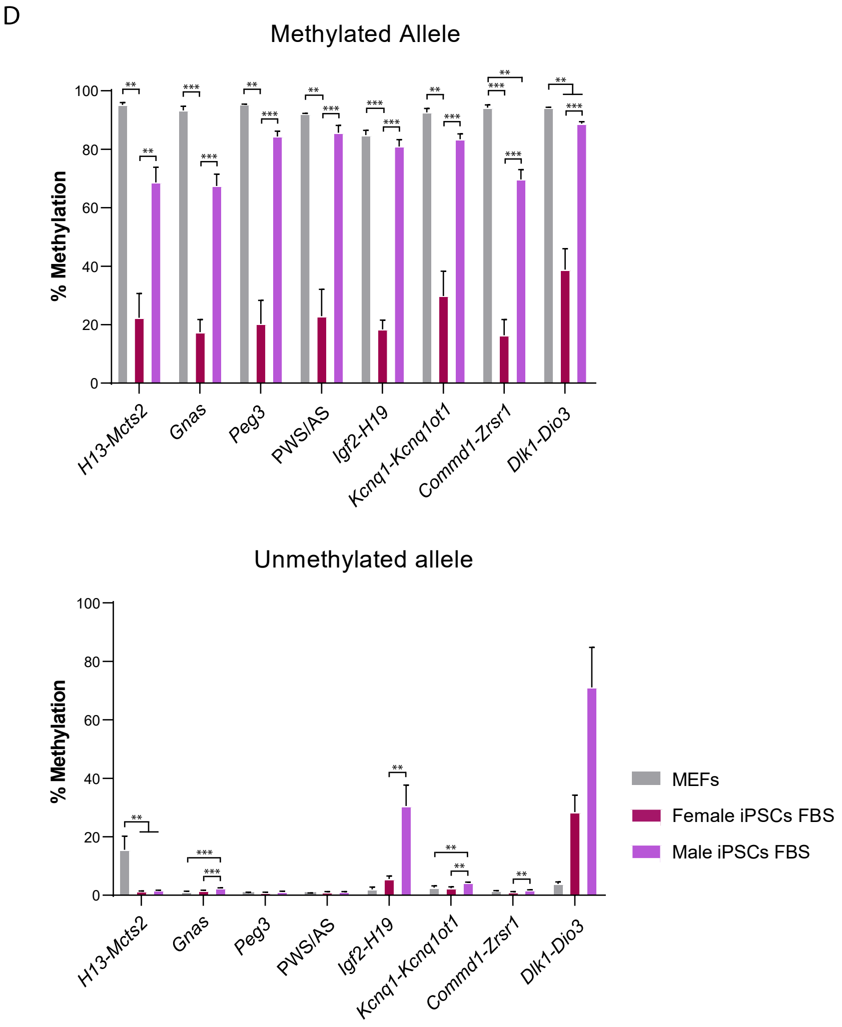


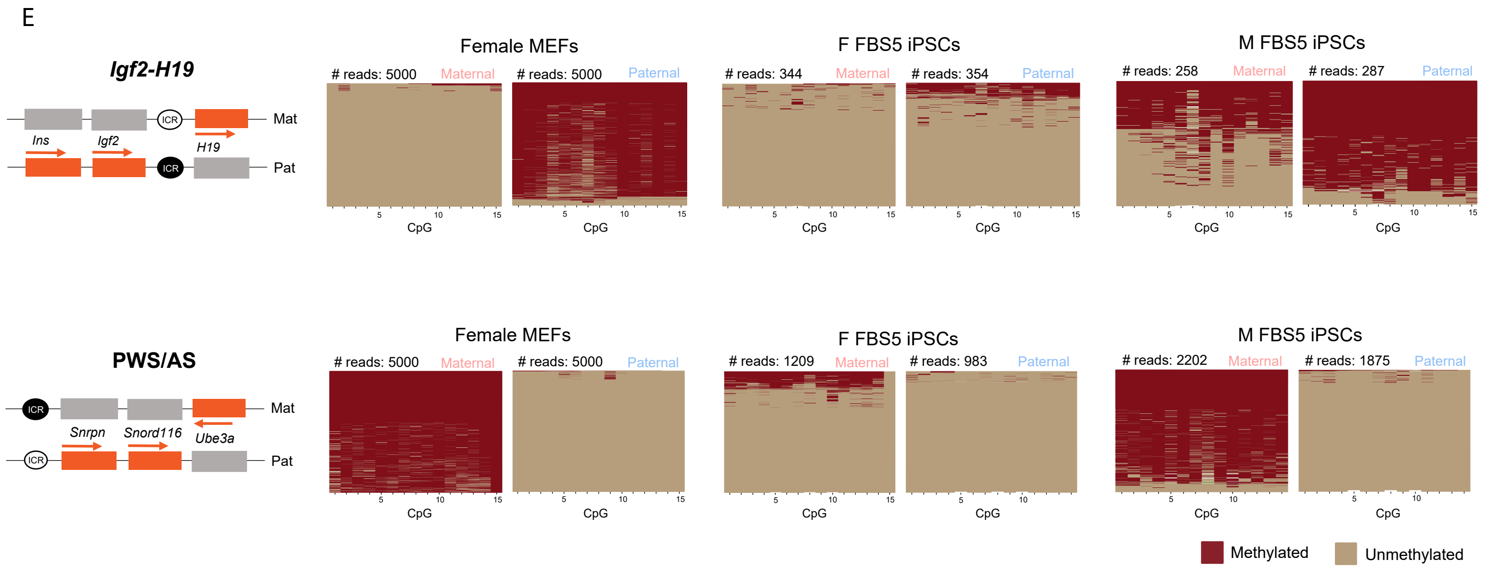

### Figure S5

A

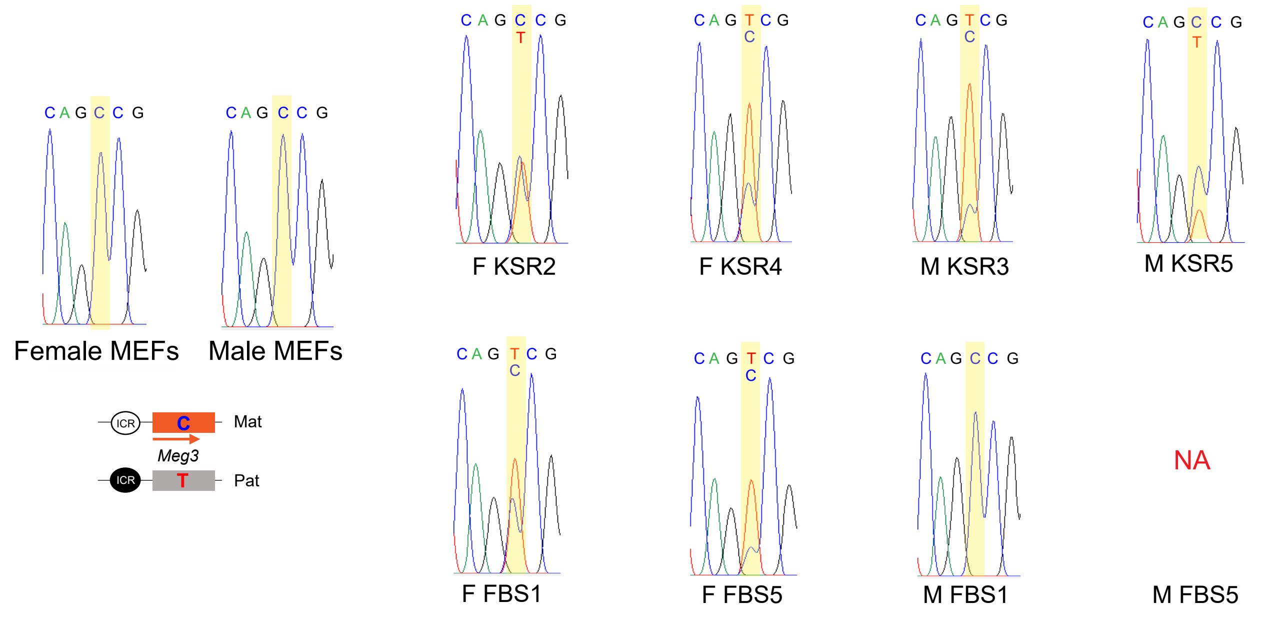

C A G C C G
C A G C C G
Female MEFs
Male MEFs
ICR
C
Mat
Meg3
ICR
T
Pat
C A G C C G
T
F KSR2
C A G T C G
C
F KSR4
C A G T C G
C
M KSR3
C A G C C G
T
M KSR5
C A G T C G
C
F FBS1
C A G T C G
C
F FBS5
C A G C C G
M FBS1
NA
M FBS5


B

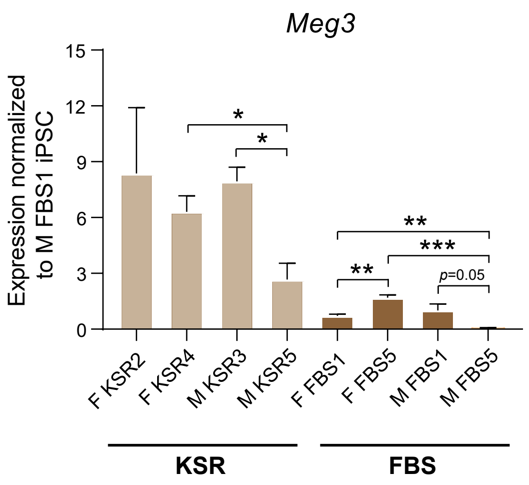

Meg3
Expression normalized to M FBS1 iPSC
15
12
9
6
3
0
*
*
**
***
**
p=0.05
F KSR2
F KSR4
M KSR3
M KSR5
F FBS1
F FBS5
M FBS1
M FBS5
KSR
FBS

### Figure S6

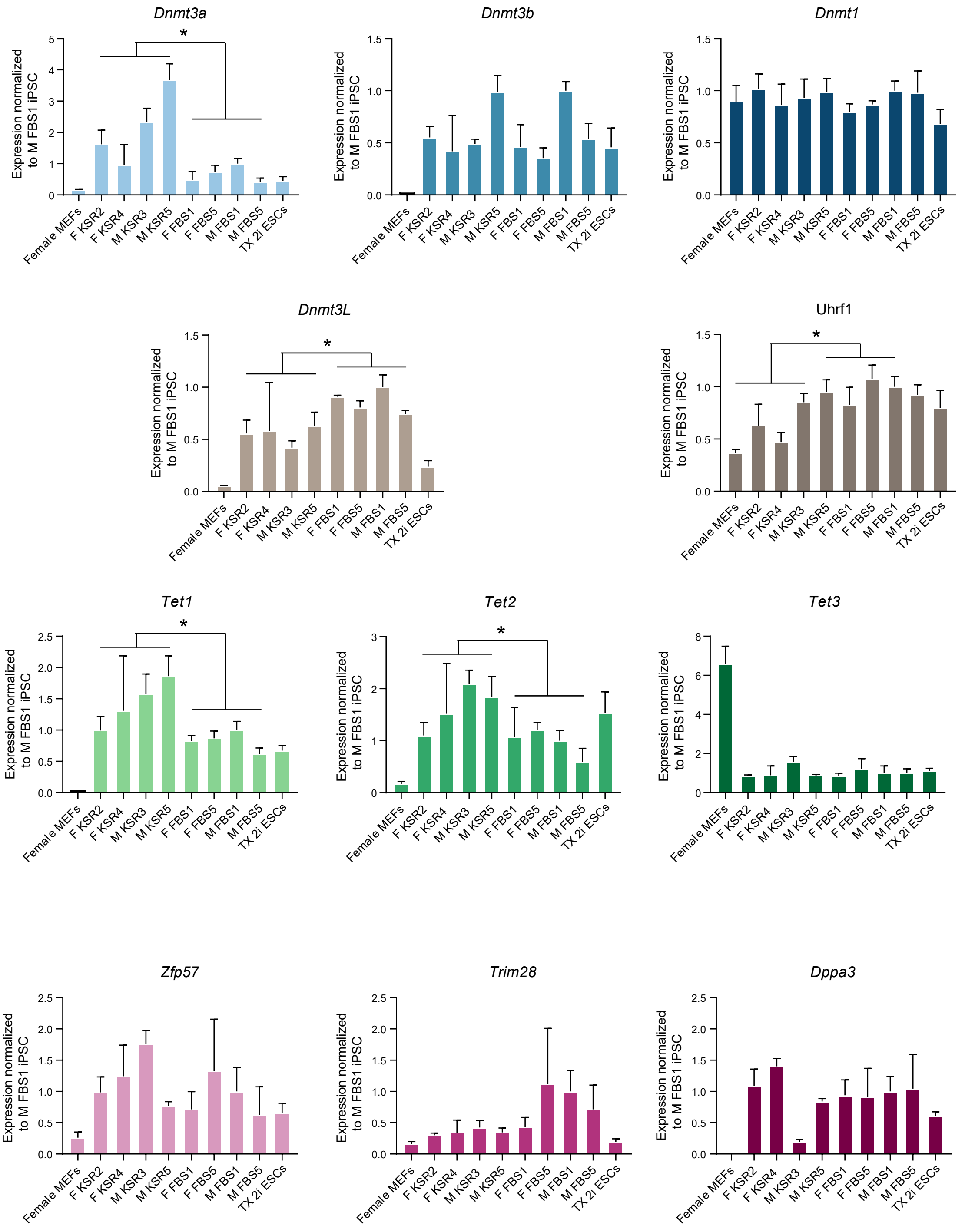

Dnmt3a
Dnmt3b
Dnmt1
Dnmt3L
Uhrf1
Tet1
Tet2
Tet3
Zfp57
Trim28
Dppa3
Expression normalized to M FBS1 iPSC
*
Female MEFs
F KSR2
F KSR4
M KSR3
M KSR5
F FBS1
F FBS5
M FBS1
M FBS5
TX 2i ESCs

**Figure S1 – Expression analysis of pluripotent markers in F1 hybrid KSR-iPSCs**

A. RT-qPCR expression analyses of pluripotent markers (*Pou5f1*, *Nanog* and *Esrrb*) normalized with the *Gapdh* housekeeping gene in female MEFs, female (F KSR1-F KSR6), male (M KSR1-M KSR6) iPSCs and JM8.F6 and TX 2i ESCs; Values for each gene were normalized to the JM8.F6 ESCs; each bar represents data from only one biological replicate.

B. Clustering analysis of the normalized RNAseq counts for all the biological triplicates of female MEFs, female (F KSR2 and F KSR4) and male (M KSR3 and M KSR5) iPSCs and TX 2i ESCs.

C. Representative IFs for three pluripotent markers (SSEA1, OCT4 and NANOG) in red and nuclei in blue (DAPI staining) in F KSR2 and M KSR5 iPSCs; Scale bars correspond to 10 µm.

D. Differential expression analysis between female and male KSR-iPSCs using both EdgeR (*p*-value < 0.05 with multiple testing correction) and intensity difference filter (*p*-value < 0.05 with multiple testing correction). Graph represents the number of upregulated and downregulated genes in female versus male KSR-iPSCs The number of differentially expressed genes from the X chromosome is highlighted. There are no downregulated X-linked genes.

**Figure S2 – Widespread imprinting methylation defects in KSR-iPSCs**

A. Methylation analysis of *Sox2* (unmethylated control) and *Prickle1* (methylated control) in male and female MEFs, female (F KSR1-F KSR6) and male (M KSR1-M KSR6) iPSCs and TX 2i ESCs; Each graph represents the mean ± SD methylation levels measured at each CpG within different genomic regions per parental allele for each sample;

B. Methylation analysis of *Gnas* and *Kcnq1-Kcnq1ot1* imprinted loci in female and male MEFs, female (F KSR1-F KSR6) and male (M KSR1-M KSR6) iPSCs and TX 2i ESCs; Each graph represents the mean ± SD methylation levels measured at each CpG within different genomic regions per parental allele for each sample;

C. Plots display methylated and unmethylated CpGs for each CpG position (in columns) in all the individual reads (in rows) for the *Gnas* and *Kcnq1-Kcnq1ot1* imprinted loci in female MEFs, F KSR2 and M KSR5 iPSCs and TX 2i ESCs.

**Figure S3 - RNAseq data in KSR-iPSCs**

A. Heatmap representing the percentage of total reads of each parental chromosome in biological triplicates of female MEFs, female (F KSR2 and F KSR4) male (M KSR3 and M KSR5) iPSCs and TX 2i ESCs.

B. Genome browser view of chromosome 6 region containing *Mest* imprinted gene (denoted by black arrow); Height of bars correspond to log2 RPKM values for each gene on either maternal (MAT) or paternal (PAT) inherited allele for biological triplicates of female MEFs, female (F KSR2 and F KSR4) male (M KSR3 and M KSR5) iPSCs and TX 2i ESCs. Selected genes names on sense (red) and antisense (dark blue) are shown.

**Figure S4 – Generation of F1 hybrid FBS-iPSCs**

A. White-field microscopy showing F FBS5 and M FBS2 iPSCs with typical morphology; scale bar represents 70 µm.

B. RT-qPCR expression analyses of pluripotent markers (*Pou5f1*, *Nanog* and *Esrrb*) normalized to the *Gapdh* housekeeping gene in female MEFs, female (F FBS1-F FBS5), male (M FBS1-M FBS5) iPSCs and JM8.F6 ESCs; Values for each gene were normalized to the JM8.F6 ESCs; each bar represents data from only one biological replicate.

C. Table and representative H&E staining of teratomas after subcutaneous injection of $2 \times 10^6$ cells into the flanks of NSG mice. In vivo iPSCs efficiently contribute to ectoderm, mesoderm, endoderm and occasionally trophectoderm. **i**, Low magnification of a mature teratoma, scale bar represents 250 µm. **ii**, Trophectoderm-derived trophoblast giant cells, associated with large vascular spaces (black arrowhead), characteristic of placental tissue. **iii**, Ectodermal components corresponding to squamous epithelium (black arrowhead). **iv**, Endodermal components corresponding to ciliated respiratory epithelium (black arrowhead). **v, vi, vii**, Mesodermal components (black arrowhead) corresponding to fibrous tissue, cartilage, and blood vessels, respectively; ii-vii scale bar represents 100 µm. Table summarizes the successful generation of teratomas with tissues from the three germ layers from F FBS1, M FBS1 and M FBS5 iPSCs.

D. Average percentage of methylation at methylated and unmethylated ICRs in both parental MEFs (note: same data as in Fig. 2B), female and male FBS-iPSCs; Graph represents the mean ± SEM methylation levels measured at each CpG within different genomic regions per parental allele for each group of samples. Statistically significant differences are indicated as ** $p < 0.01$; *** $p < 0.001$ (unpaired two-tailed Student's *t*-test).

E. Plots displaying methylated and unmethylated CpGs for each CpG position (in columns) in all the individual reads (in rows) for both the *Igf2-H19* and PWS/AS loci in female MEFs, F FBS5 and M FBS5 iPSCs; Schemes on the left of the plots represent the normal methylation status of each ICR in the *Igf2-H19* (top) and PWS/AS (bottom) imprinted regions (white circle – unmethylated ICR; black circle – methylated ICR; Mat – maternal allele; Pat – paternal allele; orange rectangles – expressed genes; grey rectangles – silenced genes; regions are not drawn to scale.

**Figure S5 - *Meg3* expression in female and make KSR- and FBS-iPSCs**

A. Allelic-specific *Meg3* expression analysis assayed by Sanger sequencing. Chromatograms are shown for female and male MEFs, F KSR2, F KSR4, M KSR3, M KSR5, F FBS1, F FBS5, M FBS1 iPSCs. M FBS5 does not express *Meg3*, hence allelic expression was not performed (NA - not applicable); schemes on the bottom of MEFs chromatograms represent the normal *Meg3* imprinting with the associated SNP of each allele; orange rectangle – maternally *Meg3* expressed gene; grey rectangle – paternally silenced Meg3 gene; region is not drawn to scale.

B. RT-qPCR expression analyses for *Meg3* normalized with the *Gapdh* housekeeping gene in F KSR2, F KSR4, M KSR3, M KSR5, F FBS1, F FBS5, M FBS1 and M FBS5 iPSCs (n=3; except for F KSR4 iPSC line, where n=2). Values for each gene were normalized to the M FBS1 iPSC. Statistically significant differences between samples (compared within medium formulation) are indicated as * $p < 0.01$, ** $p < 0.001$ and *** $p < 0.0001$ (unpaired two-tailed Student's *t*-test).

**Figure S6 – Expression analysis of genes involved in the DNA methylation machinery and imprinting protection**

RT-qPCR expression analyses for *Dnmt3a*, *Dnmt3b*, *Dnmt1*, *Dnmt3L*, *Uhrf1*, *Tet1*, *Tet2*, *Tet3*, *Zfp57*, *Trim28* and *Dppa3* normalized with the *Gapdh* housekeeping gene in female MEFs, F KSR2, F KSR4, M KSR3, M KSR5, F FBS1, F FBS5, M FBS1, M FBS5 iPSCs (n=3; except for F KSR4 iPSC, where n =2). Values for each gene were normalized to the M FBS1 iPSC. Statistically significant differences between KSR- and FBS-iPSCs are indicated as * $p < 0.05$ (two-way ANOVA followed by Tukey's multiple comparisons test).

| | Article | Stadtfeld et al., (2010) | Sun, et al., (2012) | Takikawa et al., (2013) | Yagi,et al., (2019) |
|---|---|---|---|---|---|
| Reprogramming | System | DOX-inducible, OKSM in *Col1a1* locus + *rtTA* in *Rosa26* locus | Retrovirus vsv-pseudotyped expressing individual human K,O,S,M | DOX-inducible, OSKM in *Col1a1* locus + *rtTA* in *Rosa26* locus | *PiggyBac* vector containing a DOX-inducible OSKM + *rtTA* |
| | Culture Conditions | 15% FBS | 20% KSR | 15% FBS | 15% FBS |
| iPSCs lines | Strain | ———— | Reciprocal crosses of C57BL/6J x Cast/Ei | (F)DBA/2 x (M)129 | (F)129X1/SvJ x (M) MSM/Ms |
| | Number | 6 | 8 | 12 | 5 (and 1 pre-iPSC) |
| | Gender | Not defined | Female and Male | Not defined | Male |
| Imprinting analyses | Imprinted clusters analysed | *Dlk1-Dio3* | PWS/AS, *Peg3*, *Kcqn1-Kcnq1ot1* , *Dlk1-Dio3*, *Igf2-H19*, *Mcts2-H13* | *Dlk1-Dio3*, *Igf2-H19, Rasgrf1*, *Mest/Peg1*, *Peg3*, *Sgce-Peg10*, PWS/AS, *Plagl1/Zac1* | Virtually all |
| | Methylation method | Bisulfite Pyrosequencing | Bisulfite Pyrosequencing | COBRA / Bisulfite sequencing | Bisulfite sequencing MethylC-seq |
| | Expression | Affymetrix mRNA expression microarray & RT-qPCR | Pyrosequencing | Allele-specific RT-PCR | RNA-seq |
| | Parental allele distinction | No | No | Yes, only for *Zim1* and *Snrpn* genes | Yes, large number of genes with SNPs |
| | Output | Hypermethylation of *Dlk1-Dio3* | Hypomethylation of PWS/AS, *Peg3* & *Kcnq1-Kcnq1ot1* (only female); *Dlk1-Dio3*, *Mcts2-H13* and *Igf2-H19* (male + female) | Hypomethylation of *Dlk1-Dio3*, *Igf2-H19, Rasgrf1*, *Peg3*, *Sgce-Peg10*, PWS/AS, *Plagl1/Zac1* Hypermethylation of *Mest/Peg1* | Hypermethylation of *Dlk1-Dio3* and *Igf2-H19* |

**Table S1 –** Previous studies addressing imprinting defects in mouse iPSCs

Abbreviations: DOX - Doxycycline; OSKM – Oct4, Sox2, Klf4, c-Myc; FBS – Fetal bovine serum; KSR – Knockout serum replacement; F - Female; M – Male.
